# Sensory receptor expansion and neural accommodation in butterfly color vision

**DOI:** 10.1101/2025.10.30.685642

**Authors:** Ke Gao, Julia Ainsworth, Antoine Donati, Yunchong Zhao, Michelle Franc Ragsac, Cara Genduso, Zoie Andre, Andrew Tomlinson, Michael W. Perry

## Abstract

The evolution of complex brains required existing neurons and neural circuits to accommodate new inputs. The genetic and developmental mechanisms that enable such integration are largely unknown. Butterflies evolved more complex retinal mosaics through the addition of a second R7 color photoreceptor per ommatidium (unit eye). In *Drosophila*, the unique R7 makes a stochastic choice to express one of two opsin genes. In butterflies, the two R7s make independent stochastic cell fate choices in each ommatidium, producing three ommatidial types instead of two. Here, we investigate the developmental basis of this change and how the butterfly brain accommodates expanded sensory receptor input. We first identified the changes in gene expression that cause a second R7 cell to be specified. We then modified *Drosophila* retinas to have butterfly-like transcription factor expression, causing recruitment of an additional R7. The two R7s make independent stochastic choices, like butterflies, leading to three stochastically distributed ommatidial types. In *Drosophila*, the two R7 subtypes connect to their target neurons, either yDm8 or pDm8. Dm8 neurons of both types are born in excess and Dm8s that do not find connections with their cognate y or pR7s undergo apoptosis. In the presence of extra R7s in butterfly-like fly retinas, additional Dm8s are retained, leading to two Dm8s per medulla column that make appropriate connections with the matching R7 subtypes, facilitating the expansion of color vision. We propose that the presence of cells that would otherwise die provide developmental flexibility that can allow brains to accommodate newly evolved inputs.

## Main

As animal brains evolve increased complexity, new neurons must integrate into established networks without disrupting preexisting neural function. Such a process must modify existing circuits, genes, and developmental mechanisms to accommodate new inputs without adversely impacting fitness^1–4^. Understanding how such changes have occurred is essential for uncovering the mechanisms that enable innovation in brains and sensory systems across species.

Neurons are often initially generated in excess during development. The neurotrophic hypothesis explains this process as a competition for limited survival signals: only neurons that form appropriate synaptic connections receive sufficient support, while others are eliminated^5,6^. Such overproduction may also facilitate evolutionary change by enabling new inputs to be matched with a flexible pool of potential partners^7^. Whether such mechanisms have been used during the integration of additional neurons during evolution has not been explored.

The insect visual system serves as a complex yet tractable model for the study of neural development and evolution. Like other insects, *Drosophila melanogaster* compound eyes are composed of individual repeating unit eyes (ommatidia). Each ommatidium contains eight photoreceptors (PRs, R1-8). *Drosophila* retinas contain two subtypes of ommatidia that are distributed randomly across the eye in a 70/30 ratio. The decision controlling which ommatidial subtype is produced is controlled by a probabilistic decision to express the transcription factor Spineless (Ss) in each R7 PR, yielding a mosaic of 70% Ss-ON and 30% Ss-OFF fates^8,9^. The Ss outcome controls which Rhodopsin (Rh) protein is expressed in R7, enabling color comparisons and color vision (reviewed in^10^). Ommatidia with Ss-ON R7s are called “yellow” ommatidia because of the color of the R7 rhabdom viewed using transmitted light^11^ and express Rh4 in yellow R7 (yR7) and Rh6 in the matching yellow R8 (yR8), where R8 fate is coordinated with the Ss ON/OFF decision in R7 by Activin/BMP signaling^12^. Ommatidia with Ss-OFF R7s are called “pale” and express Rh3 in pale R7 (pR7) and Rh5 in pale R8 (pR8)^10^.

Butterflies often rely on color vision for finding mates and nectar. A key evolutionary innovation in butterfly ommatidia is that they contain nine PRs instead of the ancestral eight and include two R7 PRs^13–15^. This results in two independent, probabilistic decisions to express Ss, producing ommatidia that can be ON/OFF, OFF/OFF, and ON/ON for Ss expression. The two R7s express Rhs sensitive to Blue (Ss^ON^) or UV (Ss^OFF^) light in three combinations in each ommatidium: B/UV (type I), UV/UV (type II), and B/B (type III), respectively^14^. This change in TF expression is responsible for novel eye patterning, providing butterflies with additional PR subtypes. These PRs have subsequently evolved to express new Rhs, thereby enabling expanded color comparisons^16^. The increase in butterfly retina complexity raises two questions: how are the additional R7s incorporated into ommatidia, and how are their neural projections integrated into preexisting circuits in the optic lobe?

## A common gene regulatory network

In *Drosophila*, PR recruitment occurs sequentially during the development of each ommatidium: first R8, then additional PRs are recruited as immediate neighbors of R8: R2/5, R3/4, R1/6, and finally R7 (reviewed in^17^). Three signaling pathways coordinate R7 specification: EGFR is activated by Spitz from R2/5, Notch by Delta from R1/6, and the RTK Sevenless by Boss from R8^18,19^. The order of PR recruitment is highly conserved across the insects^15^, but how this process has been modified to recruit a second R7 in butterflies is unknown.

To identify genes involved in PR specification and the recruitment of the extra R7s in butterfly ommatidia, we performed single nucleus RNA sequencing (snRNAseq) on developing retinas of the butterfly *Vanessa cardui* at 20% pupation (P20), capturing all cell types in the developing retina (Fig. 1a). We compared these data to previously published single cell RNA sequencing (scRNAseq) data from developing L3-stage *Drosophila* retinas (Fig. 1b)^20^. We identified the specific cell types corresponding to each cluster using combinations of markers that are well established in *Drosophila* and highly conserved across the insects, including Spalt (Sal) for R7 and R8, Prospero (Pros) for R7, Defective proventriculus (Dve) for “outer” PRs R1-6, and Senseless (Sens) for R8^14,15^. Additional genes that we observed in homologous cell types between *Drosophila* and butterflies included Sevenup (Svp) in R3/4/1/6, Orthodenticle (Otd/Oc) and Elav in all PRs, Rhomboid (Rho) and Sevenless (Sev) in differentiating PRs, and Cut (Ct) and Shaven/dPax2 (Sv) in cone cells (Fig. 1c and Extended data Fig. 1&2). The conserved expression of these TFs and signaling pathway members in homologous retina cell types suggested they are components of the “insect eye ground plan”^15^.

**Fig. 1.**
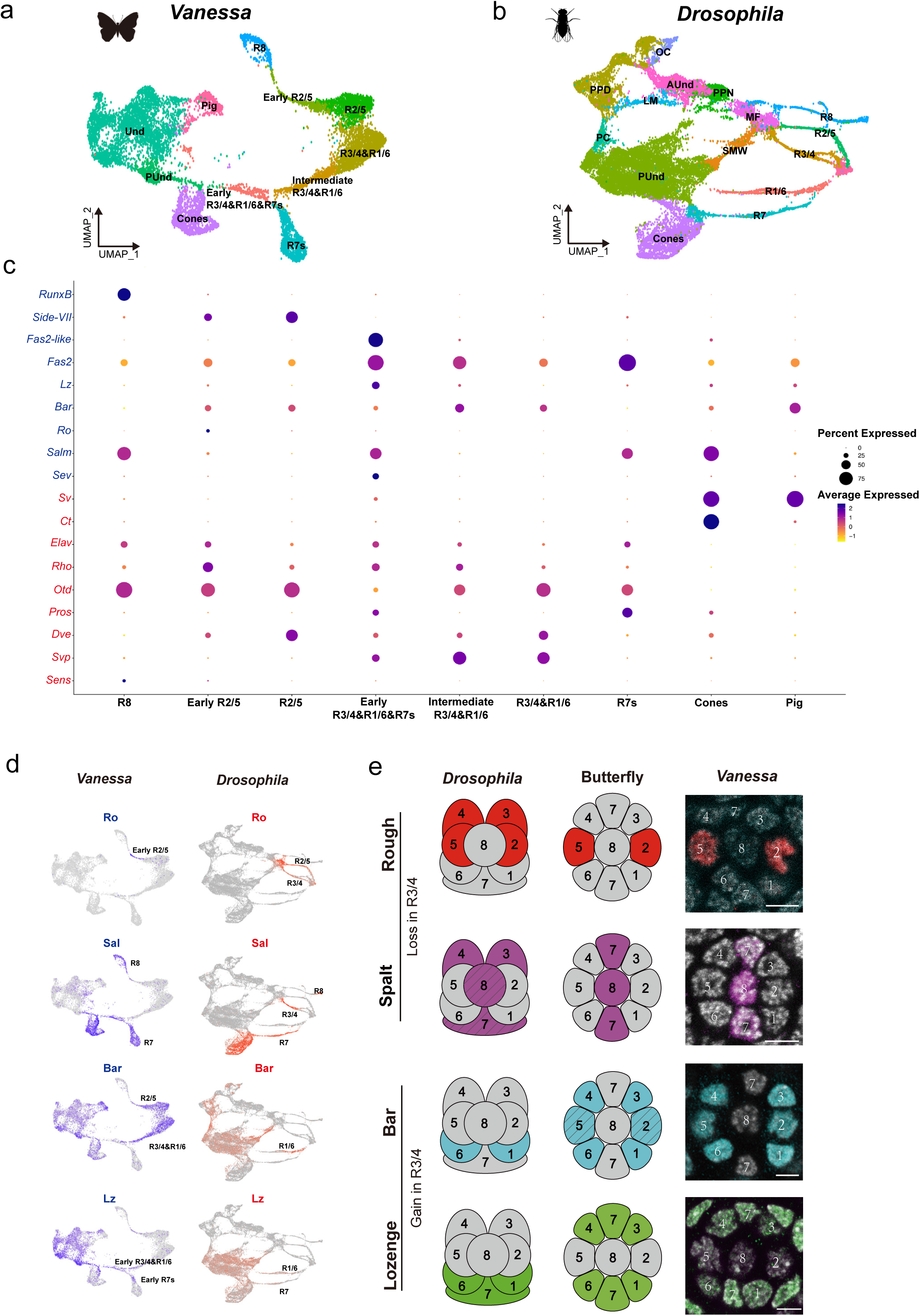
snRNAseq comparison reveals differential TF expression in the developing retina between *Drosophila* and butterfly. **a,b,** UMAP plot showing PR clusters from snRNAseq data of the butterfly P20 pupal retina (**a**), and the late larval eye disc in *Drosophila* (**b**). **c**, Dot plot showing TF expression in the butterfly retina. TFs in blue are differentially expressed between *Drosophila* and butterfly, while TFs in red are deeply conserved in both species. **d**, Feature plots highlighting four key TFs expressed in PRs between *Drosophila* and butterfly. **e**, Left: schematic representation of the expression patterns of Rough (red), Spalt (magenta), Bar (cyan) and Lozenge (green) in the *Drosophila* and butterfly ommatidium. R1-6 are outer PRs; R7/8 are inner PRs. In *Drosophila*, Rough is expressed in R2/5&R3/4, but is lost from R3/4 in butterfly. Spalt is expressed in R3/4 near the MF and later in R7/8 (shown with slashes) in *Drosophila*, but is lost in R3/4 in butterfly. Bar is expressed in R1/6 in *Drosophila*, but is gained in R3/4 and later in R2/5 (slashes) in butterfly. Lozenge is expressed in R1/6 & R7 in *Drosophila*, but is gained in R3/4 in butterfly. Right: Antibody stains show the expression of Rough, Spalt, Bar and Lozenge in a single ommatidium of the pupal retina in the butterfly *V. cardui*. Scale bars, 5 µm. Und, undifferentiated; PUnd, posterior undifferentiated; Pig, pigment; OC, ocelli; PPD, anterior peripodial; PC, posterior cuboidal margin peripodium, LM, lateral margin peripodium; AUnd, anterior undifferentiated; MF, morphogenetic furrow; SMW, second mitotic wave.

We also identified genes that were expressed differently between flies and butterflies (Fig. 1c and Extended Data Fig. 3). While Runt is coexpressed with Ss and UV-sensitive Rh4 in *Drosophila* R7 PRs^21^, Runt expression was lost in butterfly PRs (Extended Data Fig. 3). Runt paralog RunxB was instead gained in butterfly R8 (Extended Data Fig. 3a). A copy of Fasciclin-2-like (Fas2-like) that is not present in the *Drosophila* genome was expressed primarily during early PR differentiation in *Vanessa* (Extended Data Fig.1). We also observed R2/5-specific expression of Side-VII, which has not been reported in *Drosophila*. We next used HCR *in situ* hybridization to validate the expression patterns of a subset of these genes, and found that they match our snRNAseq results (Extended Data Fig. 3).

Strikingly, the cluster corresponding to the *Drosophila* R3/4 cell type is absent in *Vanessa*. Distinct clusters corresponding to R2/5, R3/4, and R1/6 PRs that are recruited as pairs during development can be identified in *Drosophila*^20^ (Fig.1b). In contrast, the UMAP plot for *Vanessa* shows a distinct R2/5 cluster and a combined R3/4/1/6 cluster approximately twice the size of the R2/5 cluster (Extended data Fig.1a). This relative proportion suggested that cells in the R3/4 position differentiate as R1/6-type in the butterfly.

To determine whether cells in the R3/4 position gain R1/6-like gene expression, we identified genes which are differentially expressed in these pairs in *Drosophila* and evaluated their expression in *Vanessa* using immunohistochemistry (Fig.1). In *Drosophila*, R3/4 are defined by their expression of the TFs Rough (Ro) and Sal, while R1/6 instead express Lozenge (Lz) and BarH1 (Fig.1). To evaluate expression in *Vanessa*, we used an existing cross-reactive antibody for Sal and generated new butterfly-specific antibodies for Ro, Lz, and Bar homologs (see Methods). In *Vanessa*, cells in the R3/4 position adopt R1/6-like fate (Fig.1e and Extended Data Fig. 4), expressing Lz and Bar but not Ro or Sal (Extended Data Fig. 4a-c), resulting in symmetric patterns of expression in each ommatidium. This confirms that cells in the R3/4 position adopt R1/6-like identity in butterflies.

The conversion of R3/4 cells to R1/6 identity suggested an intriguing hypothesis: that cells in the R3/4 position in butterflies turn on the developmental program normally used by R1/6 to recruit a neighboring cell to become R7. In *Drosophila*, when the R3/4 pair are incorporated into the ommatidium, they adhere strongly and prevent cells between them from gaining contact with the central R8^22,23^. R1/6, however, remain separate from each other allowing another cell (the R7 precursor) to lie between them and contact R8. In developing *Drosophila* ommatidia, a cell initially lies between R3/4s but is excluded from contacting R8 as R3/4 move closer together. This is the ‘mystery cell’ (M cell), which subsequently regresses from the cluster and rejoins the surrounding cell pool^22,23^. We propose that the M cell is the precursor of the supernumerary R7 in the butterfly, and that the respecification of R3/4 as R1/6 prevents them from zipping up, allowing the cell to retain contact with R8 and differentiate as an R7 PR.

We set out to determine which specific changes in gene expression are sufficient for the conversion in R3/4 to R1/6 fate, and whether that conversion is sufficient to allow recruitment of a second, butterfly-like R7 in the position of the mystery cell, opposite the existing R7.

## Recruitment of the additional R7 requires Lz expression in R3/4

While the regulatory relationships between TFs expressed in R3/4 or R1/6 had been examined previously^24–29^, these studies did not focus on whether changes in R3/4 fate could induce recruitment of an adjacent R7, and this therefore became our objective. We first conducted gene loss-of-function experiments and examined the expression of Lz and Bar as markers of R1/6 fate, and Sal and Ro as markers of R3/4 fate.

We made MARCM clones in *Drosophila* eye discs using null alleles of *ro* and *sal* (Fig. 2). In Ro loss-of-function clones (marked by GFP in Fig. 2b-e), GFP-marked cells in the R3/4 position do not lose Sal expression (Fig. 2c) and do not gain Bar (Fig. 2d) or Lz expression (Fig. 2e). This indicated that Ro expression in R3/4 is not required for Sal activation or for repression of Bar or Lz. In Sal loss-of-function clones (Fig. 2g-j), GFP-marked cells in the R3/4 position do not lose Ro (Fig. 2h) or gain Bar (Fig. 2i) or Lz (Fig. 2j), indicated that Sal was also not required for Ro activation, or for repression of Bar or Lz. This suggested that neither Ro nor Sal are fully upstream of each other, Bar, or Lz, and their removal is not sufficient for the recruitment of a neighboring R7.

**Fig. 2.**
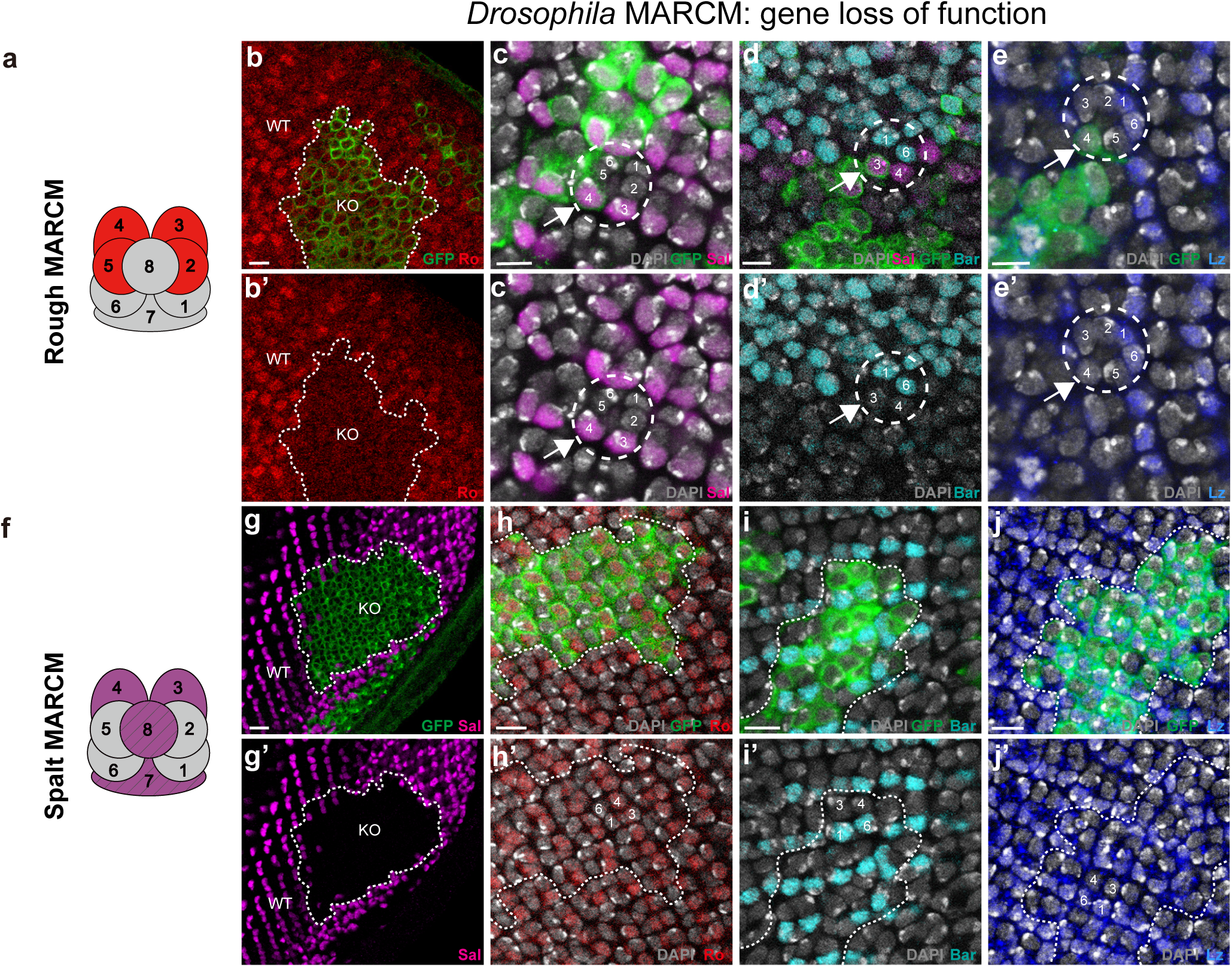
Rough and Spalt loss of function in *Drosophila* does not de-repress Lozenge or Bar. **a**, Schematic showing Rough expression in R2/5, R3/4 in *Drosophila* L3 larval eye discs. **b,b’**, Rough (red) is lost in MARCM mutant clones (KO regions labelled by GFP, green), as compared to wild type (WT) region. Scale bars, 5 µm. **c-e’,** loss of Rough does not affect Spalt, Bar and Lozenge expression. The circle in (**c,c’**) highlights an ommatidium where Rough (GFP, green) is lost in R4 (arrows), while Spalt expression in R4 (magenta) is unchanged. The circle in (**d,d’**) highlights an ommatidium where Rough is lost in R3 (arrows), and Bar is not gained in R3/4 (Spalt-positive cells in magenta), and normally expressed in R1/6 (cyan). The circle in (**e,e’**) highlights an ommatidium where Rough is lost in R4 (arrows), Lozenge is not gained in R3/4, and normally expressed in R1/6 (blue). Scale bars, 5 µm. **f**, schematic showing Spalt expression in R3/4, R7/8 in *Drosophila*. **g-g’**, Spalt (magenta) is lost in MARCM mutant clones (KO regions labelled by GFP, green), compare to WT region. Scale bars, 10 µm. **h-j’**. loss of Spalt does not affect Rough (red in **h,h’**), Bar (cyan in **i,i’**), and Lozenge (blue in **j,j’**) expression. An ommatidium labelled with PR numbers in white is shown in the Spalt MARCM KO regions. Scale bars, 5 µm.

We then used CRISPR/Cas9 in *Vanessa* to knock out *bar* and *lz*, producing mosaic mutant animals. We used immunohistochemistry to identify regions of tissue where the targeted protein is lost. In Bar loss-of-function clones, expression of Ro, Sal, and Lz was unaffected (Fig. 3c-j and Extended Data Fig. 5a-e). In Lz loss-of-function clones, however, both R7 PRs are lost as indicated by loss of Pros and Sal expressing cells flanking the Sal-positive R8 (Fig. 3l-o). When examining the boundaries of Lz knockout clones, we observed examples of ommatidia where Lz positive outer PRs (R1/6 type) are adjacent to a Lz-negative clone and have a missing R7, indicating that Lz expression was also required autonomously in R7 (Extended Data Fig. 5f,g) (as in *Drosophila*)^26^. Within null clones, the number of PRs around each R8 is variable from 6 to 8, suggesting that R7s could either be variably lost or transformed into outer PRs, as observed in *Drosophila lz* mutant ommatidia^24,25^. Outer PRs within Lz mutant regions have variable levels of Bar and Ro expression (Fig. 3p-s), suggesting a derepression of Ro in the absence of Lz. In general, we observed an inverse relationship between Ro and Bar expression, suggesting that Ro represses Bar in *Vanessa* (Fig. 3q-s). Overall, these results demonstrate that Lz is necessary for the recruitment of both R7s, and suggest that Ro, Bar, and Sal are downstream of Lz expression in defining R1/6 fate.

**Fig. 3.**
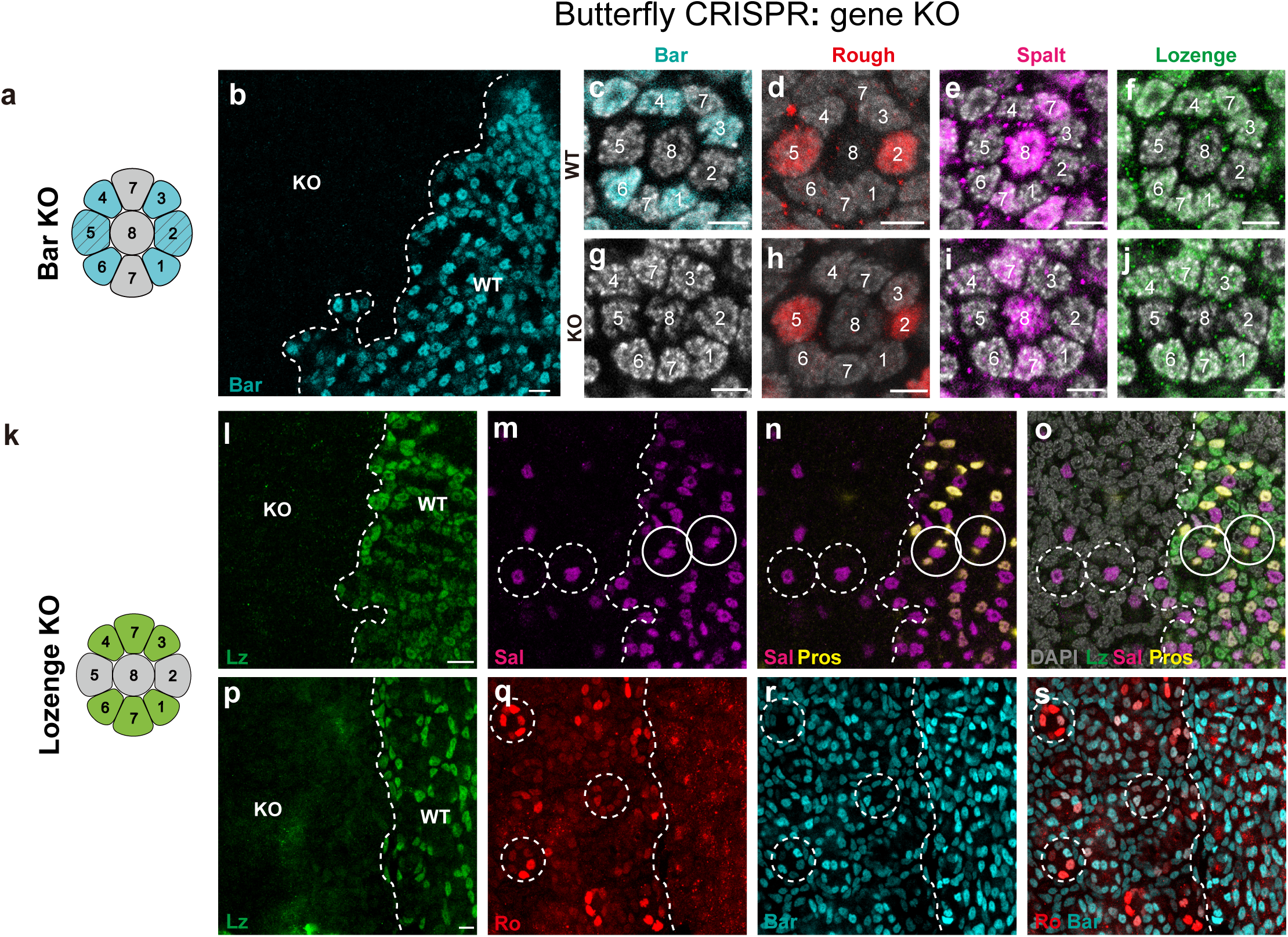
Lozenge is necessary for the recruitment of two R7s in butterflies. **a**, Schematic showing Bar expression in R1-6 in a butterfly ommatidium. **b-j**, CRISPR mosaic knockout of Bar does not affect the expression of Rough, Spalt or Lozenge in the pupal retina of *V. cardui*. **b**, Bar is lost in KO regions compared to WT region (cyan). The boundary between WT and Bar KO regions is indicated by a white dashed line. Scale bars, 10 µm. **c-f,** a single ommatidium in the WT region. **g,** a single ommatidium in the Bar KO region, where Rough (red, **h**), Spalt (magenta, **i**) and Lozenge (green, **j**) are normally expressed. Scale bars, 5 µm. **k**, schematic showing Lozenge expression in R1/6, R3/4, and R7s in butterfly. **i-o**, CRISPR mosaic knockouts of Lozenge result in the loss of R7s. Dashed circles highlight example ommatidia in Lozenge KO region, as compared to solid circles in WT regions. The boundary between WT and lozenge KO regions is indicated by a white dashed line. In the WT region, Spalt (magenta) is expressed in R8 and two R7s per ommatidium, while Prospero (yellow) is expressed in two R7s. In KO regions, Prospero and Spalt coexpressing R7 cells are lost, but not Spalt-only expression cells (R8s). Scale bars, 10 µm. **p-s,** In Lozenge KO regions, we observe a variable number of outer PRs (dashed circles), indicating that R7s can be variably lost or transformed into outer PRs. Outer PRs within KO regions show mixed identity, with variable levels of Rough (red in **q**) and Bar (cyan in **r**) expression, suggesting derepression of Rough in the absence of Lozenge. Scale bars, 10 µm.

We next sought to add Lz to R3/4 cells in *Drosophila* to determine whether this change is sufficient to induce recruitment of a neighboring R7. A similar experiment had been performed previously with an unexpected result: R3 and R4 themselves were variably transformed into R7^30^, and it was determined that Notch levels are important in whether they are converted. R1/6 express Lz and have low Notch (N) activity, while R3/4 lack Lz and have high N signaling. N signaling in R3/4 is critical for coordinated ommatidial rotation in *Drosophila,* but potentially less important in butterflies, which have fused rhabdoms and lack neural superposition^31^. We thus aimed to ectopically express Lz in R3/4 while simultaneously reducing N activity to determine whether cells in the R3/4 position would become R1/6-like and allow the incorporation of the M cell as an R7 PR.

We tested a number of drivers (see Supplementary Table 4) and found that supplying Lz to R3/4 using sev.lz^30^ while reducing N activity using sev.Su(H)enR^19^ produced butterfly-like ommatidia. We identified ommatidia near the morphogenetic furrow at L3 that had nine PRs with two R7s in the same position as in butterflies, in the position of the M cell (Fig.4b). This fate becomes especially clear at pupal stages (Fig.4c). This phenotype was not fully penetrant, with some *Drosophila*-like ommatidia remaining, and rare ommatidia had two extra R7s at the position of the mystery cell (Fig.4c). This experiment succeeded in recruiting M as an R7 and produced butterfly-like ommatidia.

By pupal stages (P50), these butterfly-like ommatidia made a stochastic, cell intrinsic choice to express Ss: we observed ommatidia with either two, one, or no Ss-positive R7s (Fig. 4f, dashed circles), in contrast to wild type that had ommatidia that were either Ss-ON or Ss-OFF in the single R7 (Fig. 4e, dashed circles). Finally, we examined Rh expression downstream of Ss in adult retinas and found ommatidia that have two R7s which resemble the butterfly-like three-way stochastic choices, i.e., two yR7s, two pR7s, or one yR7 and one pR7 (Fig. 4d). Therefore, when the mystery cell is recruited as an additional R7 PR, it leads to three stochastically distributed ommatidial types instead of two. Together, these results demonstrate that conversion of R3/4 toward R1/6 fate is sufficient to implement the developmental program used to recruit a neighboring R7 PR.

**Fig. 4.**
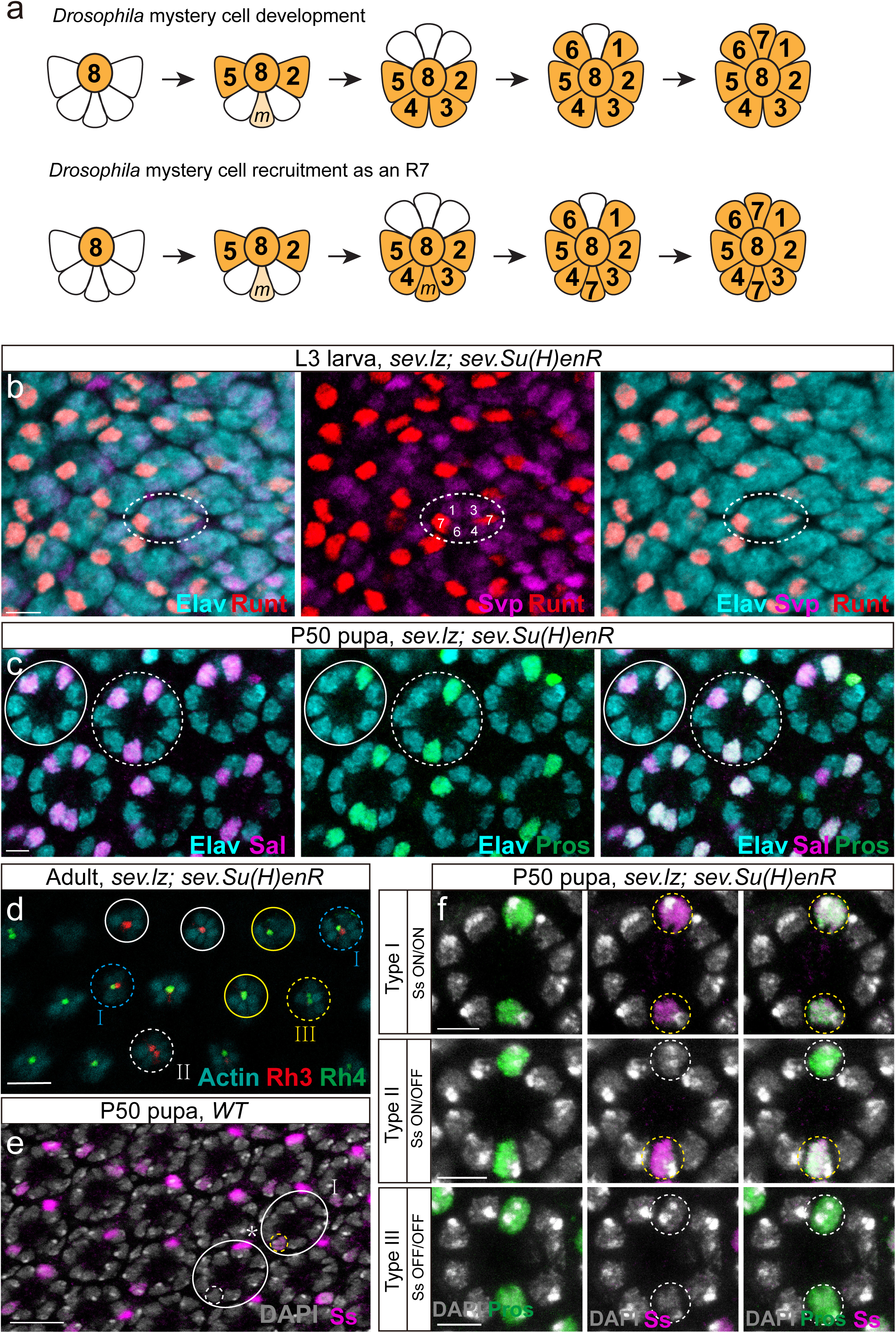
Conversion of R3/4 toward R1/6 fate is sufficient to create butterfly-like ommatidia in *Drosophila*. **a**, Schematics show the order of PR recruitment in *Drosophila* eye imaginal disc. The mystery cell(s) do not present after R2/5 differentiation in WT flies (top), but the mystery cell(s) recruit as R7s in flies with the genotype sev.lz, sev.Su(H)enR flies (bottom). **b,** Third-instar larval eye disc with genotype sev.lz, sev.Su(H)enR. Ommatidia (Elav labeling PRs in cyan) are observed with two R7s (Runt-positive, red), while R3/4&R1/6 are labelled by Svp (magenta). Scale bars, 5 µm. **c**, P50 retina with genotype sev.lz, sev.Su(H)enR. Ommatidia are observed with two R7s (Spalt in magenta and Prospero in green; coexpression in white; Elav in cyan). A solid circle shows a WT ommatidium, while a dotted circle highlights an ommatidium with an additional R7. Scale bars, 5 µm. **d**, Adult retina of sev.lz, sev.Su(H)enR. Ommatidia appear butterfly-like, with either two yR7s expressing Rh4 (green) (III, yellow dashed circles), two pR7s expressing Rh3 (red) (II, white dashed circles), or one yR7 and one pR7 expressing Rh3 and Rh4 respectively (I, blue dashed circles). WT ommatidia show one yR7 (yellow solid circles), and one pR7 (white solid circles). Scale bars, 10 µm. **e**, In the P50 WT retina, two types of ommatidia, circled in solid white, are observed: Ss^ON^(dotted circle in yellow) and Ss^OFF^(dotted circle in white). Scale bars, 10 µm. **f**, In P50 retina of sev.lz, sev.Su(H)enR, three types of ommatidia are observed: Ss^ON^/ Ss^ON^(type I), Ss^ON^/ Ss^OFF^ (type II) and Ss^OFF^ / Ss^OFF^ (type III). Scale bars, 5 µm.

## How does the optic lobe accommodate two R7 cells?

We next sought to understand how input from additional R7 PRs could be accommodated at the level of the optic lobe that processes visual information. Were additional, complementary genetic changes required to integrate the additional R7s into color vision circuits?

In *Drosophila*, each R7 and R8 projects to a corresponding column in the medulla where they connect to specific cell types. The primary interacting partner of R7 PRs is Dm8^32^. All Dm8s establish primary connectivity with a single R7 axon terminal in what is referred to as that Dm8’s “home column” but also interact with multiple neighboring R7s to enable center-surround comparisons^33,34^.

Recent work identified two types of Dm8s that interact with stochastically specified R7s called “yellow” or “pale” Dm8s (yDm8 or pDm8), which interact with yR7s or pR7s, respectively ^33,34^. The two Dm8 subtypes are specified deterministically in excess and send projections to find partner R7s of the correct type^33,34^. yR7s express a cell surface protein Dpr11 which interacts with DIPγ expressed in yDm8 cells^35^. This interaction allows recognition of yR7 with its cognate yDm8 and provides a survival signal: Dm8s that do not find connections undergo apoptosis ^33,34^. A similar matching mechanism must exist for pR7 and pDm8, although the specific proteins that mediate pDm8 survival remain unidentified. This matching process ensures that cell types produced via a stochastic decision in the retina are met with the appropriate types of target cells that are specified independently and deterministically.

Both types of Dm8s are produced in an excess of ∼30%^34^. We propose that this surplus allows the system to accommodate population-level variation in the stochastic ratio of R7 subtypes. While the population averages 70% yR7s, the stochastic ratio in the retina ranges from 19% to 83% Ss-ON in inbred lines^36^, demonstrating significant potential for variability. In this scenario, individuals that inherit higher or lower R7 subtype ratios produce enough excess Dm8s of each type to accommodate variation in the stochastic ratio. Such variation could provide a reason for selection to maintain an excess pool of potentially interacting Dm8s of each type.

We hypothesized that an evolutionary change in the retina that increases the number of R7s could be immediately accommodated by the extra Dm8s. To test this model, we assessed whether butterflies have two Dm8 home column projections in each medulla column, and then whether *Drosophila* modified to produce additional R7s can accommodate this input by retaining additional Dm8s.

Fig. 5a shows a *Drosophila* medulla with R7 and R8 PR terminals marked by Chaoptin (Chp). R7 axons terminate in the deeper M6 layer and interact with yDm8 projections which are labeled by DIPγ expression, as shown previously, as well as with unlabeled pDm8s^33,34^(Fig. 5a,b). The *Drosophila* Chp and DIPγ antibodies did not cross-react in butterflies and we made new antibodies to the butterfly proteins (see Materials and Methods). Corresponding side views of a butterfly medulla show similar co-localization of Chp and DIPγ in a single medulla layer. Unlike in *Drosophila*, however, we observed two distinct R7 axons within single medulla columns in butterflies (Fig. 5g, h-l). DIPγ signal was seen alongside the paired R7 terminals in a variable fashion: sometimes present on one side, both sides, or neither side, indicating one, two, or no yDm8s (Fig. 5e,m,n). There is no marker for pDm8s or a general marker for all Dm8s in butterflies, but the presence of two DIPγ-positive projections in single columns suggested that butterflies have two Dm8s per medulla column.

**Fig. 5.**
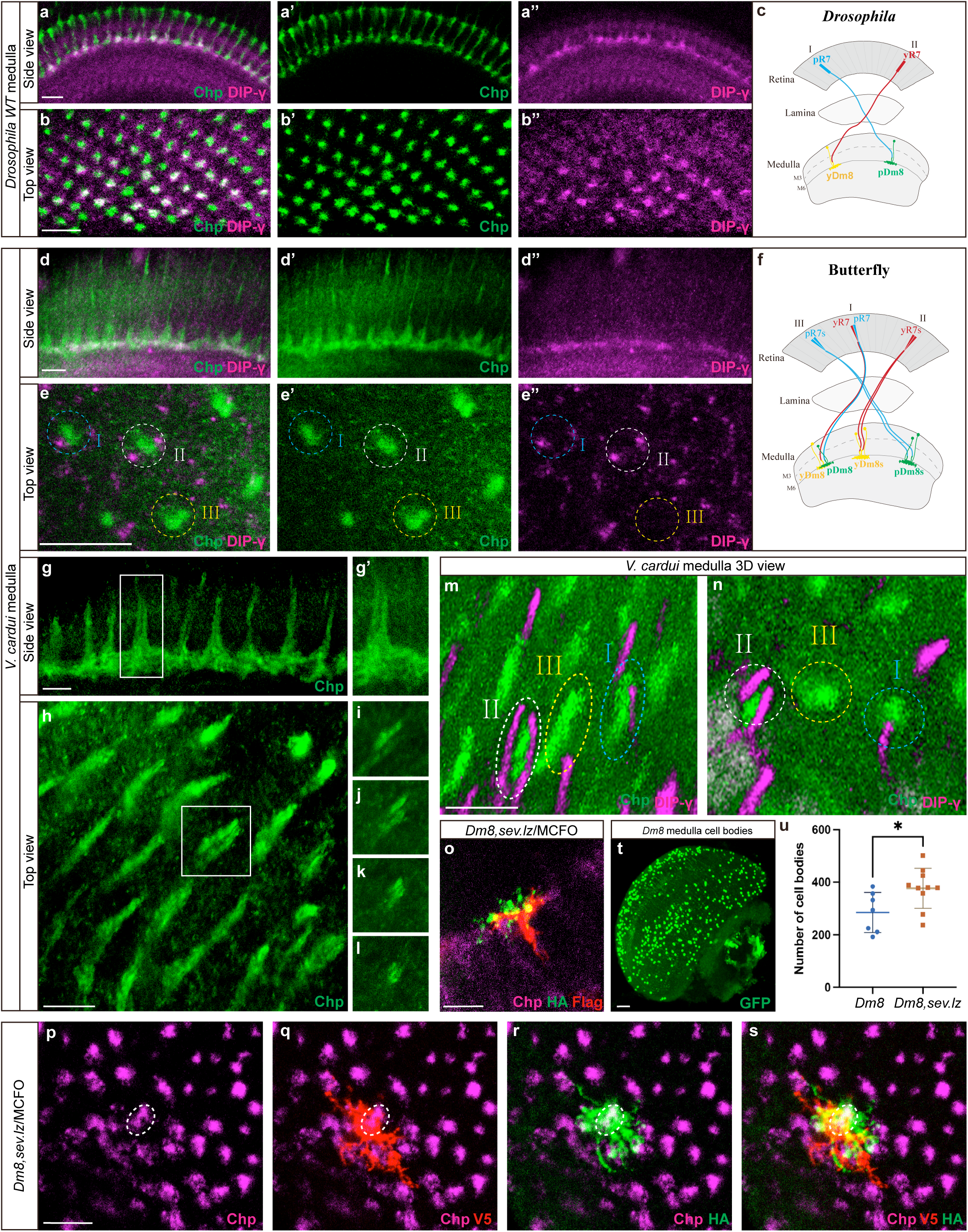
Additional R7 PRs are accommodated at the level of the brain. **a-a’**, Sideview of the P50 medulla in *Drosophila*. R7 axons are labelled by Chaoptin (green), and DIPγ (magenta) marks yDm8s. Scale bars, 10 µm. **b-b’**, Top-down view of the P50 medulla in *Drosophila*, R7 terminals are labelled by chaoptin (green), and yDm8 are marked by DIPγ (magenta). Scale bars, 5 µm. **c**, Schematic depicting the interaction of yDm8s with Ss-ON yR7s, while pDm8s interact with Ss-OFF pR7s in *Drosophila*. **d-d’**, Sideview of the adult butterfly medulla. R7 axons are labelled by chaoptin (green), and Dm8s are labelled by Dip-γ (magenta). Scale bars, 10 µm. Scale bars, 5 µm. **e-e’**,Top-down view of the adult butterfly medulla, DIPγ signal is observed alongside paired R7 terminals: present on one side (I, blue dashed circle), both sides (II, white dashed circle), or neither side (III, yellow dashed circle). Scale bars, 10 µm. **f**, Schematic showing three types of interaction between R7s and Dm8s in the butterfly. **g-i**, Two R7 axons within single columns in butterfly medulla. Scale bars, 10 µm. **m,n**, 3D views show DIPγ-positive projections along both sides of some R7 terminal pairs: one side (I, blue dashed circle), both sides (II, white dashed circle), or neither side (III, yellow dashed circle). Scale bars, 10 µm. **o-s**, MCFO labelling of two Dm8s sharing a single “home column” in sev.lz flies, which produces extra R7s per ommatidium. Two Dm8s (labeling with V5 and HA, red and green) connect to two R7 terminal axons (Chp, magenta) in one column. Scale bars, 10 µm. **t**, Dm8 cell bodies are labelled with GFP in WT. Scale bars, 20 µm. **u**, Quantification of Dm8 number in WT vs sev.lz adult medulla (WT, n=7, sev.lz, n=10). A significant difference is indicated by an asterisk (Welch’s *t* test, *P*<0.05).

We next evaluated the effect of R7 loss on Dm8 number in butterfly medulla columns. We showed previously that Lz KO causes loss of R7 PRs (Fig. 3k-o). We examined the optic lobes of Lz CRISPR mosaic somatic mutant butterflies and observed gaps in DIPγ expression in regions where R7 terminals are missing (Extended Data Fig. 8), indicating that R7 terminals are required for the retention of DIPγ-positive Dm8 neurons in retinotopic positions. This provided evidence that, as in the fly, a trophic signal from yR7s is also required for yDm8 survival in butterflies, and that DIPγ plays a conserved role in yR7/yDm8 patterning downstream of the stochastic choice in the retina.

We next examined how two R7 in each *Drosophila* ommatidium impact Dm8 retention. We used MCFO sparse labeling^37^ to label a small number of Dm8 neurons per medulla in the sev.lz background^30^, which produces extra R7s per ommatidium (Fig. 5o-s). When extra R7s are present, we identified cases where two Dm8s share a single home column, which we did not observe in WT controls and which has not been observed in previous studies of Dm8 morphology of WT animals examined using either light microscopy^33,34^ or electron microscopy ^38,39^. Although the labeled Dm8s have home columns in the same position (Fig. 5o), their lateral projections to neighboring columns differ, suggesting that they have different center-surround connections. These results suggested that two Dm8s can occupy the same home column when additional R7s are present. We next compared the overall number of Dm8s retained to adulthood in each background. We quantified the number of Dm8s present in WT vs sev.lz adult brains (Fig. 5t) and observed a 32% increase per brain when additional R7s were present (Fig. 5u, Welch’s *t* test, df=15, *P*=0.027).

Finally, we examined whether *Drosophila* that produce butterfly-like ommatidial types are able to achieve proper pairing between their two R7s and Dm8 subtypes within single medulla columns. Fig. 6a-c shows pairing in WT control medullas, with DIPγ-GFP in yDm8 projections matching Rh4-lacZ/βGal-positive yR7 terminals. GMR-RFP labels all R7 terminals. We crossed the same reporters into the butterfly-like sev.lz, sev.Su(H)enR background and unlike WT, observed examples of two R7 axons within the same medulla column (Fig. 6d-i). In medulla columns containing two Rh4-lacZ-positive yR7s, home column projections from two DIPγ-positive yDm8 neurons are present (Fig. 6d,e,h and Extended Data Fig. 10a,b). In contrast, in columns containing one yR7 and one pR7, only a single DIPγ-positive yDm8 home column projection is observed and is located alongside the yR7 terminal (Fig. 6f,g,i and Extended Data Fig. 10c,d). In WT, lateral projections interact with R7 terminals in several neighboring columns, and in these examples, DIPγ signal also accumulates along the distal end of the immediately adjacent pR7. Importantly, only one home column was present in heterotypic ommatidia and it has subtype specificity. In columns containing two pR7s, we did not observe home column projections of DIPγ-positive yDm8 neurons (Extended Data Fig. 10a).

**Fig. 6.**
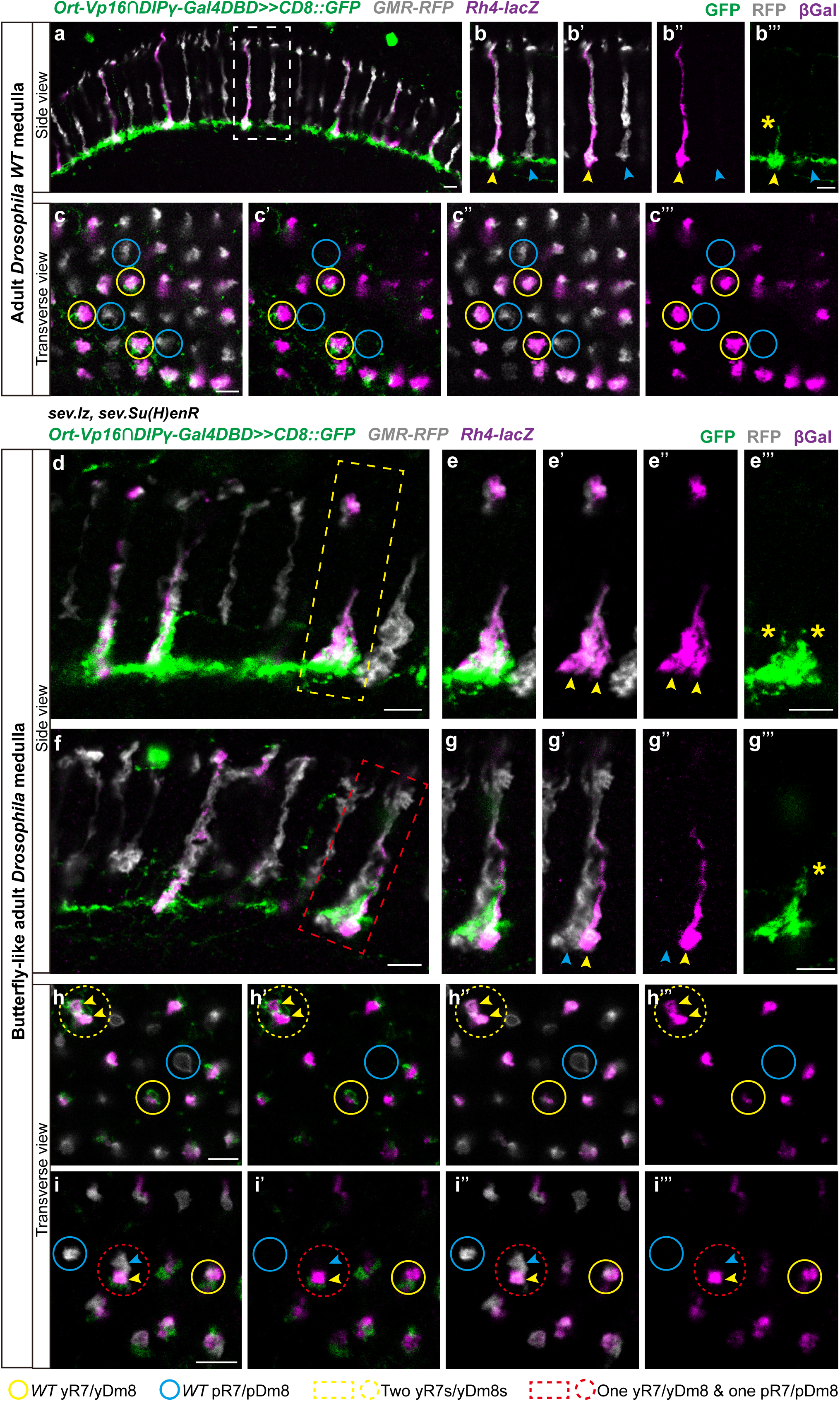
Butterfly-like *Drosophila* brains show specific connections between y/p R7s and y/p Dm8s subtypes. **a-b’’’**, side view of the adult medulla in WT. All R7 axons are labelled by GMR-RFP (gray). yR7 axons are labelled by βGalactosidase (magenta). yDm8s are labelled by DIPγ (green). Arrowheads in (**b-b’’’**) indicate the yR7 (yellow) and pR7 (blue) within a single home column in WT. The yellow asterisk in (**b’’’**) marks the yDm8 neuron interacting with the yR7 in the home column. **c**-**c’’’**, top-down view of the adult medulla in WT. R7 terminals are labelled by GMR-RFP (gray). yR7 terminals are labelled by βGalactosidase (magenta). yDm8s are labelled by DIPγ (green). Yellow circles highlight DIPγ-positive yDm8 neurons paired with yR7s, whereas blue circles indicate pR7s lacking associated DIPγ-positive yDm8 neurons. **d-g’’’**, side view of the adult medulla in sev.lz, sev.Su(H)enR flies. The yellow dashed rectangle in (**d**) highlights two yR7 axons labelled by βGalactosidase (magenta) within a single home column. Two yellow arrowheads in (**e-e’**) indicate the two yR7 axons (magenta), and the two yellow asterisks in (**e’’’**) mark two DIPγ-positive yDm8 neurons (green) within the same home column. The red dashed rectangle in (**f**) highlights two R7 axons, one yR7 labelled by βGalactosidase (magenta) and one pR7 labelled by GMR-RFP (gray) within a single home column. The yellow arrowhead in (**g’-g’’**) indicates the yR7 axon and the blue arrowhead indicates the pR7 axon, and the yellow asterisk in (**g’’’**) marks a single DIPγ-positive yDm8 neuron (green) within the same single home column. **h-i’’’**, top-down view of the adult medulla in sev.lz, sev.Su(H)enR flies. Compared with WT columns containing one yR7 (yellow solid circle) and one pR7 (blue solid circle), yellow dashed circles in (**h-h’’’**) highlight two yR7 terminals (labelled by βGalactosidase, magenta) connect to two DIPγ-positive yDm8 neurons within a single column. Two yellow arrowheads indicate the two yR7 terminals. Red dashed circles in (**i-i’’’**) highlight two R7 terminals, one yR7 labelled by βGalactosidase (magenta, yellow arrowhead) and one pR7 labelled by GMR-RFP (gray, blue arrowhead) within a single home column, and one DIPγ-positive yDm8 neuron (green) connects to yR7 terminal. Scale bar, 5 µm.

Together, this evidence suggested that butterflies produce two Dm8 neurons per medulla column and demonstrates that this evolutionary transition can be recapitulated in flies. The production of butterfly-like ommatidia in flies can immediately cause the retention of additional Dm8 neurons, and the extra R7s are immediately able to make new subtype specific connections without requiring additional genetic changes that modify the brain.

## Discussion

The developmental mechanisms that control core insect eye development are deeply conserved, despite a vast array of morphological differences and significant divergence in visual function across the insects^15^. There is evidence that the same signaling pathways are used in this patterning process and that the TFs that define PR types are conserved in species as distant as crickets^15^. The canonical eight-PR, single R7 arrangement is also highly conserved^15,40^. One of the most dramatic deviations from this conserved patterning system is found in the Lepidoptera: butterflies have increased the complexity of their stochastic retinal mosaics by producing two R7-type PRs, expanding their color vision^13–15^. Here, we identify the genetic and developmental basis of how butterflies produce two R7 PRs per ommatidium.

The most immediate benefit to having a second R7 per ommatidium should be an increase in both sensitivity and color resolution. Ommatidia that have two UV-sensitive PRs that view the same point in the visual field should have increased UV sensitivity. Similarly, Blue/Blue ommatidia would have higher sensitivity to blue light. The UV/Blue ommatidial type provides the ability to discriminate both UV and Blue light in a single position, effectively increasing color resolution compared to detecting a single wavelength range in that position. Importantly, this addition only functions effectively if each R7 in heterotypic ommatidia are able to make subtype-specific connections with the right type of Dm8. A subtler potential benefit is that while Dm8s have a central home column in the medulla that interacts with a primary R7, they also connect to a stochastic subset of neighboring ommatidial columns, enabling “center surround” comparisons. Having two R7s that each connect to a different Dm8 neuron, each with a different set of center-surround connections, could also affect color discrimination and color processing. Evolutionarily, the presence of three ommatidial types provided additional positional information and places to differentially express newly duplicated Rhs that in some cases acquired novel spectral sensitivities, such as in *Heliconius*^41^ and *Pieris*^42^ butterflies. By producing additional ommatidial types, other coordinated changes could also occur, such as the evolution of ommatidia type-specific expression of green– or red-sensitive Rhs in outer PR homologs in *Papilio* swallowtails^16^ downstream of the stochastic choices in R7^14^.

We provide evidence that insect brains are able to immediately accommodate changes in the number of R7s per ommatidium. However, there are only 30% excess of each type of Dm8 produced^33,34^, leading to a potential shortage if the entire retina were modified to have two R7s per ommatidium at once. Such a change could have initially evolved in a subregion of the retina. Supporting this idea, we identified a moth species that does just this: in *Manduca sexta* hawkmoths only the ventral retina contains ommatidia with two R7s and is butterfly-like, while ommatidia in the dorsal retina have single R7s and are fly-like^15^. A later expansion in the number of Dm8s produced could enable expansion of the region containing two R7s to the full retina, as found in butterflies.

Other studies have experimentally recapitulated events during the evolution of color vision. Some primates such as squirrel monkeys are dichromatic and lack red color vision, as do other mammals including mice. Two studies introduced human red opsin regulatory and coding sequences into PRs in either squirrel monkeys or mice and found that, amazingly, this conferred the ability to gain trichromatic vision^3,43^. The exact mechanism and molecules that mediate this gain in ability are not understood. Perhaps the expansion to trichromacy in vertebrate color vision also made use of excess neurons that are normally removed during development.

The production of more neurons than are needed is common in neural development. In some cases, the excess neurons are thought to enable neural plasticity via neural activity-dependent reinforcement of a subset^4^. The example we describe in butterfly visual systems is different: excess Dm8 neurons are produced to enable matching between a stochastically patterned retina and a deterministically patterned optic lobe, with the excess produced also able to accommodate population-level variation in the stochastic ratio. This mechanism results in developmental flexibility that offers an alternative to classic neurotrophic models by emphasizing not just cell number, but the need to match cell identity, providing a flexible framework for accommodating novel inputs. Whether excess neurons are used to accommodate stochastic patterning or to enable neural plasticity, their presence could provide potential for cooption into new roles during neural evolution.

## Methods

### Animals

Painted lady butterflies *Vanessa cardui* were obtained from Carolina Biological Supply Company, Burlington NC, USA. Larvae were fed on artificial diet individually in plastic cups. Adults were placed in a mesh cage with sugar water under artificial lights. Eggs were collected using sunflower stems. *Drosophila melanogaster* were reared on standard medium, and maintained in incubators at 25 °C.

### snRNAseq analysis

We dissected 16 pupal retinas at Day 2 from the butterfly *V. cardui*. After dissection, retinas were flash frozen in liquid nitrogen and stored at –80°C. We used 10x Multiome kit for library preparation with a target recovery of 6000 cells. Retinas were split into two biological replicates, each with a target sequencing depth of ∼300 million reads. Sample processing and sequencing were performed at the Center for Epigenomic, University of California, San Diego. snRNAseq data were processed using the Cell Ranger pipeline (v9.0.1) with the *V. cardui* reference genome ilVanCard2.1. The combined raw matrix from Cell Ranger output had 13907 cells and a total of 577 million reads. We performed downstream analyses in R (v4.4.2) using Seurat (v5.2.1). We first filtered out low quality cells, and normalized the filtered gene expression matrix using the NormalizeData function with the LogNormalize method and a scaling factor of 10000. To visualize cell clusters using UMAP, we performed dimensionality reduction with RunPCA (npcs=150), followed by clustering and visualization with RunUMAP (dims=1:50), FindNeighbors, and FindClusters (resolution=1.0) in Seurat. Differentially expressed genes for clusters were identified using the FindAllMarkers function with a minimum expression threshold (min.pct=0.05) and a log-fold change threshold (logfc.threshold=0.05). For comparison, snRNAseq data from *Drosophila* larval eye disc were used from GEO (accession no. GSE235110)^20^.

### Fly genetics

The following genotypes of *D. melanogaster* were used: **a**) Mosaic analysis with a repressible cell marker (MARCM) clones: yw122, UAS-CD8gfp;; tubgal4, frt82, tubgal80/Tm6b (BDSC #86311), yw, hsflp; sp/Cyo; rox63, FRT82/Tm2 (BDSC #6335), UAS-CD8gfp, hsflp122; frt40a, tubgal80; tubgal4/Tm6b (BDSC #44406), yw, frt40, Df(1)sal/CyO (from Claude Desplan); **b**) MultiColor FlpOut (MCFO): pBPhsFlp2:: PEST in attP3;; HA_V5_FLAG (BDSC #64093), pBPhsFlp2:: PEST in attP3;; HA_V5_FLAG_OLLAS (BDSC #64091), w;; Dm8gal4 (BDSC #49087), w;; Dm8gal4, sev.lz/Tm6b (sev.lz from Andrew Tomlinson), yw; sp/CyO; UAS-CD8GFP/Tm6b (BDSC #7465); **c**) Mystery cells: yw,hsflp;sp/CyO; sev.lz/Tm2, yw,hsflp;;sev.Su(H)enR/Tm2 (sev.Su(H)enR from Andrew Tomlinson); **d**) butterfly-like fly: GMR-myr::RFP; OrtC1-3-Vp16/UAS-CD8::GFP; sev.lz,sev.Su(H)enR/DIPγ-Gal4DBD, Rh4-lacZ.

To generate MARCM clones, fly larvae were heat-shocked at 37 °C in a water bath for 30-45 min after a 48h egg laying period^44^. To label Dm8 neurons using MCFO, newly hatched fly adults were heat-shocked at 37 °C in a water bath for 12-20 min^45^.

### Butterfly CRISPR/Cas9 knock-out

To determine the functional roles of Bar and Lozenge in the recruitment of two R7s, we knocked out *bar* and *lozenge* in *V. cardui* using CRISPR/Cas9. The details of butterfly embryo injection were described in ref.^14^, and are summarized here briefly: eggs were collected 1-7h after egg laying, two sgRNA and Cas9 were co-injected with yellow sgRNA for targeting *yellow* color mutation. After injection, the hatched larvae were immediately transferred to the artificial diet until pupation. A mixture of final concentration of sgRNA/Cas9 at 250 ng/ul was used for injection. Synthetic sgRNAs were obtained from Synthego, and sequences can be found in Supplementary Table 2. Genomic DNA was extracted from the mosaic crispants and WT butterflies using the DNeasy Blood & Tissue kit (Qiagen, Germany). PCR amplification was performed using the primers listed in Supplementary Table 1, and the PCR products were purified using the FastPure Gel DNA Extraction Mini Kit (Vazyme, China). Sanger sequencing of the sgRNA-targeted regions was conducted at Azenta Life Sciences (USA). CRISPR-induced indels (*bar* and *lz*) were confirmed using Synthego ICE (Extended Data Fig. 9).

### Immunohistochemistry and HCR *in situ* Hybridization

Larval eye/antennal imaginal disc, pupal and adult retina and optic lobes were dissected and stained as described in detail in ref.^14^. Commercial primary antibodies were used as follows: sheep anti-GFP (Bio-rad 4745-1051, 1:500), chicken anti-GFP (MilliporeSigma 06-896, 1:500), rabbit anti-mcherry (BioVision 5993-100, 1:500), mouse anti-βGalactosidase (DSHB 528101, 1:500), mouse anti-Rough (DSHB 528456, 1:10), mouse anti-svp (DSHB 2618080, 1:100), rat anti-Elav (DSHB 528217, 1:50), mouse anti-lozenge (DSHB 528346, 1:10), mouse anti-prospero (DSHB 528440, 1:10), mouse anti-chaoptin (DSHB 528161, 1:50), rabbit anti-HA (Invitrogen 26183, 1:400); sheep anti-HA (Invitrogen OSH00021W, 1:400), chicken anti-FLAG (Exalpha Biologicals 15242, 1:1000), mouse anti-V5 (Invitrogen R960-25, 1:100), rabbit anti-V5 (Invitrogen MA5-32053, 1:400). The following antibodies were gifts: guinea pig anti-SalM (1: 400), rat anti-VcBar (1:500), guinea pig anti-Ss (1:500) guinea pig anti-runt (1:1000), guinea pig anti-rh4 (1:500) (Claude Desplan, NYU).

The following polyclonal antibodies were generated for this study by Genscript (Piscataway, NJ), with sequences used for antibody production listed in Supplementary Table 2: guinea pig anti-VcDIP-γ (1:200), rabbit anti-VcRo (1:100), guinea pig anti-VcLz (1:100), rabbit anti-VcChaoptin (1:50), and we re-made a cross-reactive rabbit anti-Sal that had stopped working (1:100) (originally from^14,46^). AlexaFlour secondary antibodies were used as follows: donkey anti-rabbit-405 (1:500), donkey anti-rabbit-488 (1:500), donkey anti-mouse-488 (1:500), donkey anti-rat-488 (1:500), donkey anti-sheep-488 (1:500), donkey anti-mouse-555 (1:500), donkey anti-guinea pig-555 (1:500), donkey anti-rat-555 (1:500), donkey anti-chicken-555 (1:500), donkey anti-rat-647 (1:500), anti-guinea pig-647 (1:500), donkey anti-rabbit-647 (1:500), donkey anti-mouse-647 (1:500), phalloidin (Invitrogen A12379, 1:250).

Sequences used for hybridization chain reaction (HCR) are listed in Supplementary Table 3. HCR probes were designed as described in ref.^47^.

Images of immunohistochemical stains were acquired using a Leica SP8 confocal microscope. Confocal image stacks were processed with ImageJ and Adobe Photoshop (v23.4.1). All figures were made in Adobe IIIustrator (v29.5.1).

### Statistical analysis

The difference of the number of Dm8s present between WT and sev.lz adult brains was compared by two-sided, unpaired Welch’s *t* test (data fit assumptions of normality) in Fig. 5u. Data are shown as mean ± s.d. with individual values. Statistical significance was shown as *P*<0.05. Data analyses and plot were made in Prism (v10.4.2).

## Accession codes

### Primary accessions

#### Sequence Read Archive

SRPxxxxxx – Data will be deposited upon or before manuscript acceptance

### Data deposits

Data will be deposited upon or before manuscript acceptance at SRA under accession code SRPxxxxxx.

## Acknowledgements

We thank Claude Desplan, Noah Rose, and Emma Farley for feedback and discussion. We thank Emily Troemel and the UCSD School of Medicine Microscopy Core for the use of confocal microscopes and support (grant P30 NS047101). J.A. was supported by grant GM133351 from the NIH. This project was supported by a Hellman Fellowship (Hellman Fellows Fund, San Francisco, CA), NSF CAREER award 2339031 to M.W.P., and NIH NHGRI grant R01HG013634 to M.W.P from the EDGE program.

## Author information

### Authors and Affiliations

**Department of Cell & Developmental Biology, University of California San Diego, La Jolla, 92093, California, USA**

Ke Gao, Julia Ainsworth, Antoine Donati, Yunchong Zhao, Zoie Andre, & Michael Perry

**Halicioğlu Data Science Institute, University of California San Diego, La Jolla, CA, 92093, USA**

Michelle Franc Ragsac

**Department of Biology, New York University, New York, 10003, New York, USA**

Cara Genduso

**Department of Genetics and Development, College of Physicians and Surgeons, Columbia University, New York, NY, 10032, USA**

Andrew Tomlinson

**Zuckerman Mind Brain Behavior Institute, Columbia University, New York, NY, 10027, USA**

Andrew Tomlinson

### Contributions

K.G. and M.W.P. designed research; K.G., J.A., Y.Z., C.G., Z.A., and M.W.P. performed experiments; K.G., A.D., M.F.R., A.T., and M.W.P. analyzed data; and K.G. and M.W.P. wrote the manuscript.

### Corresponding author

Correspondence and requests for materials should be addressed to Michael Perry.

## Ethics declarations

### Competing interests

The authors declare no competing financial interests.

## Extended data figures and tables

**Extended Data Fig. 1.**
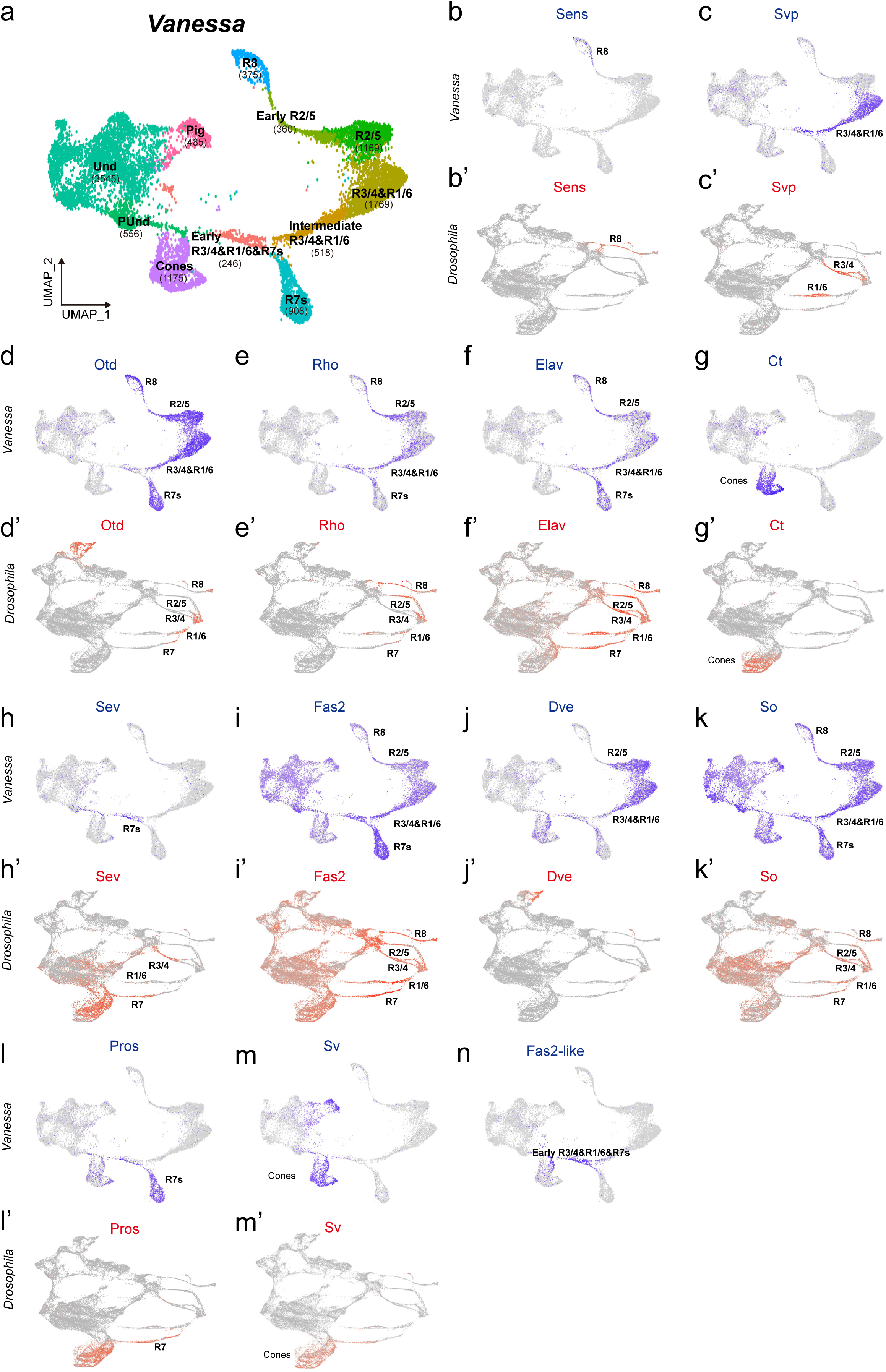
FeaturePlots highlighting TF expression in the cell types of Drosophila *and* Vanessa. **a**, UMAP plot showing clusters with a total of 11,106 cells from the snRNAseq data of the butterfly P20 pupal retina. The number of cells in each cluster is shown in brackets. Early R3/4/1/6 + Intermediate R3/4/1/6 = 764, Early R2/5 = 360, **b-n**, FeaturePlots showing the expression of TFs in PRs of *Drosophila* and *Vanessa*. Und, undifferentiated; PUnd, posterior undifferentiated; Pig, pigment.

**Extended Data Fig. 2.**
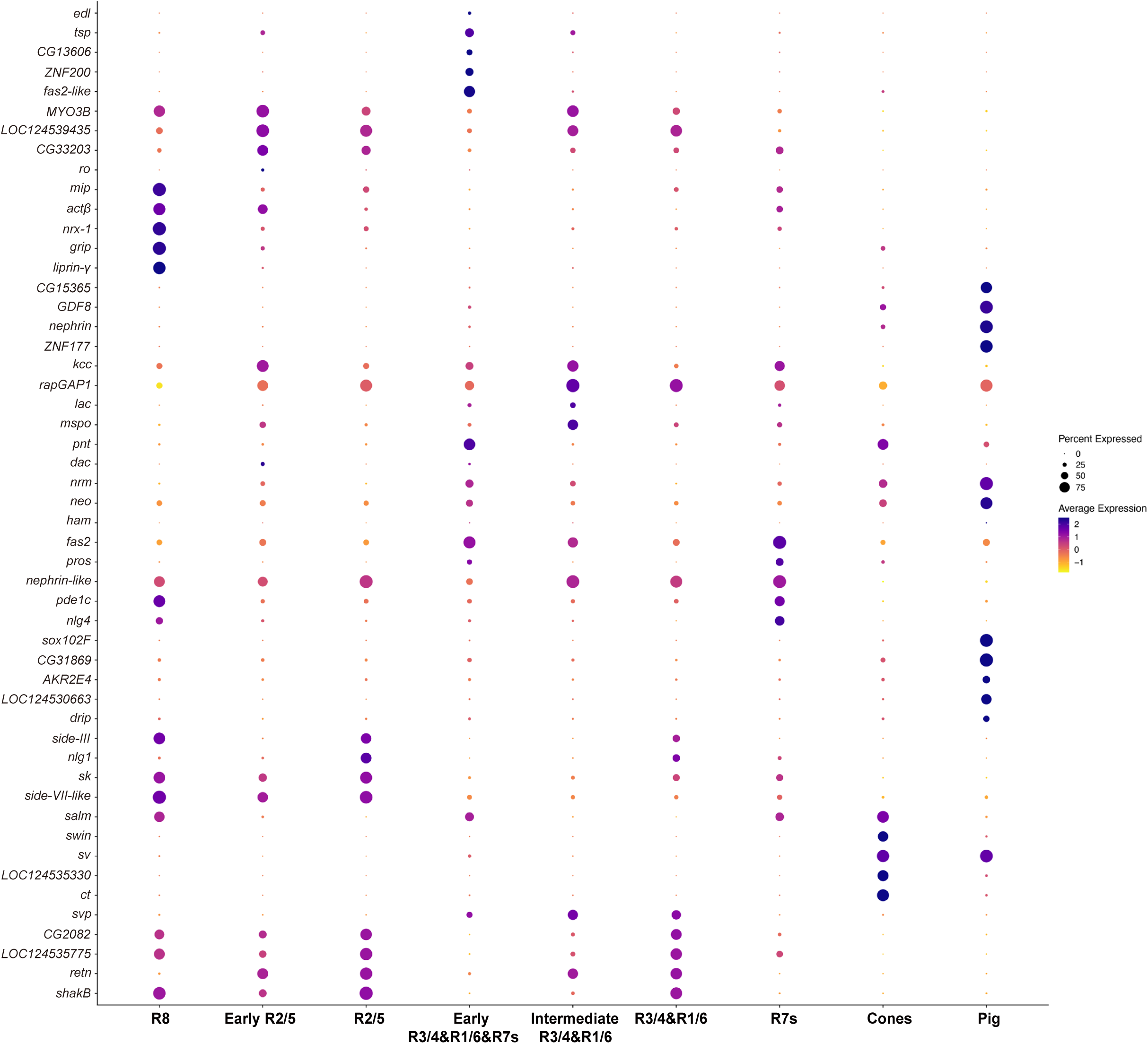
Marker gene expression within cell types. DotPlot showing the top 51 genes associated with each cell type cluster.

**Extended Data Fig. 3.**
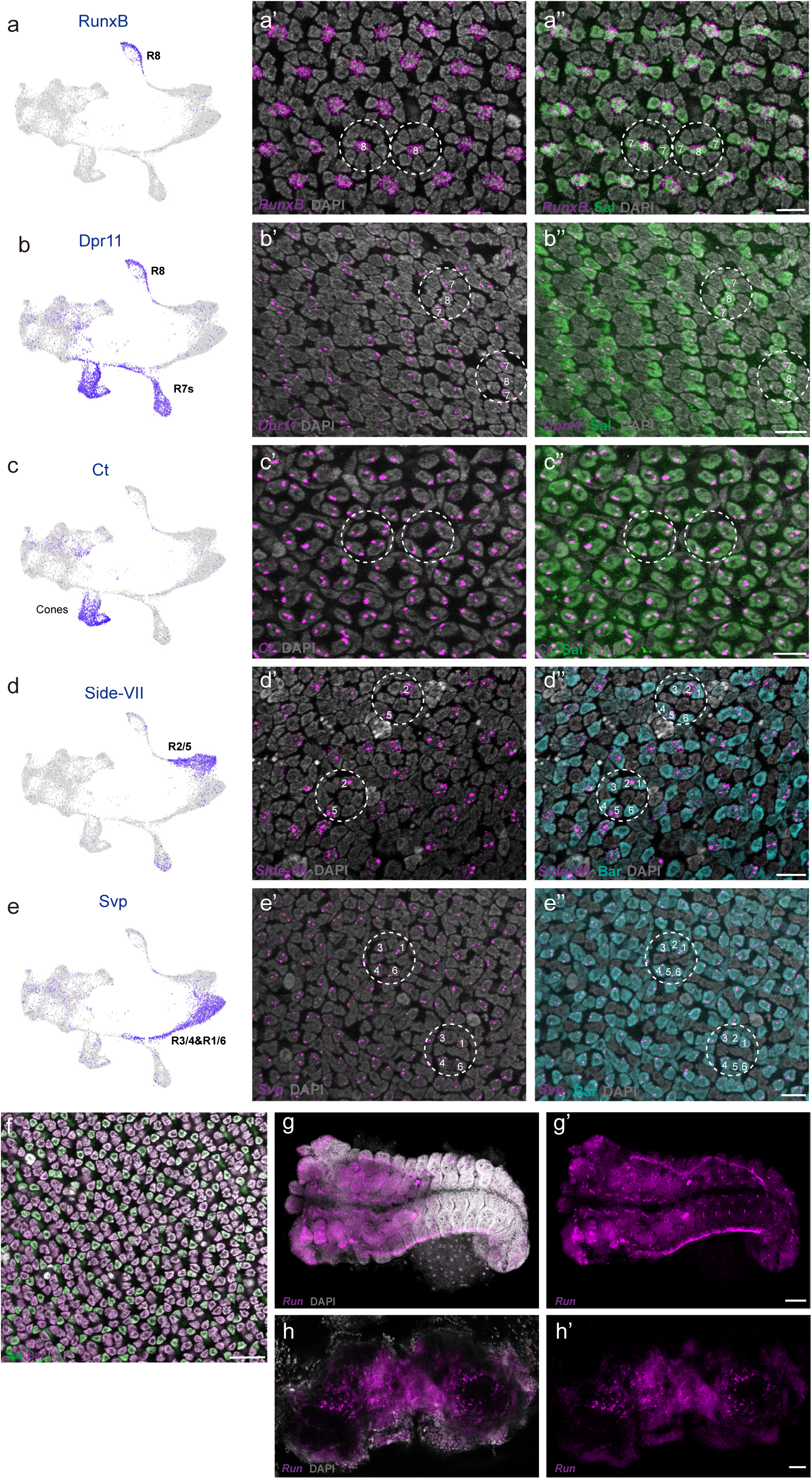
Validation of cell type clusters using HCR *in situ* hybridization and antibody staining in *Vanessa*. **a-a’**’, RunxB is expressed in R8 cells. FeaturePlot showing *RunxB* expression in R8 cluster (**a**). HCR *in situ* hybridization combined with antibody staining shows *RunxB* mRNA expression (magenta) in R8 (**a’**), Sal (green) labels R8 and R7s (**a’’**). **b-b’**’, Dpr11 is expressed in R8 and R7 cells. FeaturePlot showing *Dpr11* expression in R8 and R7s cluster (**b**). HCR *in situ* hybridization combined with antibody staining shows *Dpr11* mRNA expression (magenta) in R8 and R7s (**b’**), Sal (green) labels R8 and R7s (**b’’**). **c-c’**’, Ct is expressed in cone cells. FeaturePlot showing *Ct* expression in cone cells cluster (**c**). HCR *in situ* hybridization combined with antibody staining shows *Ct* mRNA expression (magenta) in cone cells (**c’**), Sal (green) labels cone cells (**c’’**). **d-d’**’, Side-VII is expressed in R2/5 cells. FeaturePlot showing *Side-VII* expression in R2/5 cells cluster (**d**). HCR *in situ* hybridization combined with antibody staining shows *Side-VII* mRNA expression (magenta) in R2/5 (**d’**), Bar labels outer PRs, R1-6 (cyan) (**d’’**). **e-e’**’, Svp is expressed in R3/4 and R1/6 cells. FeaturePlot showing *Svp* expression in R3/4&R1/6 cells cluster (**e**). HCR *in situ* hybridization combined with antibody staining shows *Svp* mRNA expression (magenta) in R3/4 and R1/6 (**e’**), Bar (cyan) labels outer PRs, R1-6 (**e’’**). **a’**-**e’’**, scale bar 10µm. **f**, *Runt* mRNA expression is not detected by HCR *in situ* hybridization combined with antibody staining, only Sal (red) and Bar (cyan) expression is observed. Scale bar 20µm. **g-h’**, *Runt* mRNA expression (magenta) is observed in the embryo (**g**,**g’**) and adult brain (**h**,**h’**). **g**-**g’**, scale bar 100µm. **h**-**h’**, scale bar 50µm. All nuclei are labeled with DAPI (gray).

**Extended Data Fig. 4.**
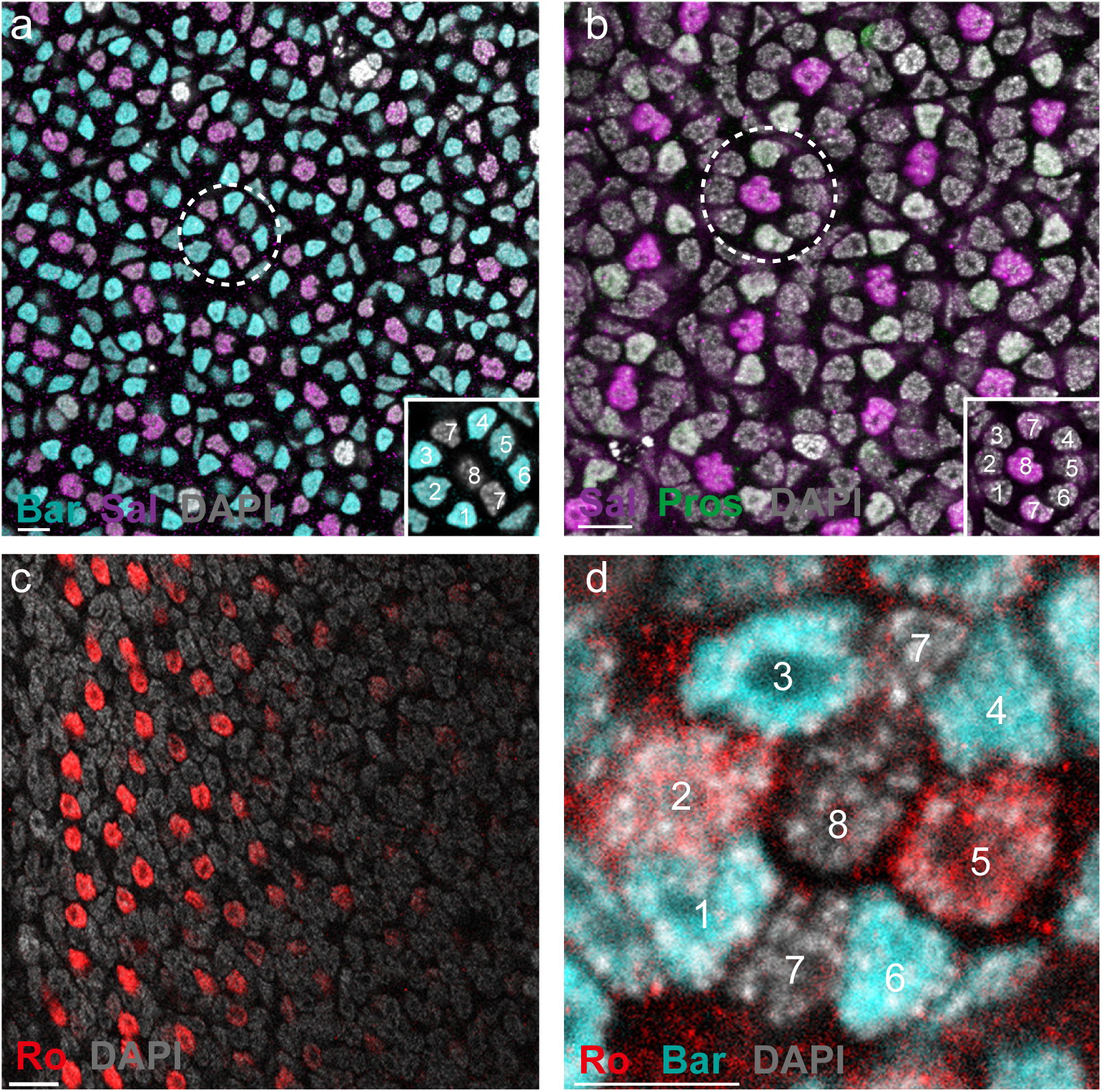
Expression of four TFs in butterfly PRs. **a**, Antibody stains showing ommatidia in the butterfly *Vanessa*. Bar (cyan) is expressed in outer PRs, R1-6. An ommatidium with nine PRs is shown within a dashed circle (also see inset). Scale bar, 10µm. **b**, Sal (magenta) is expressed in R8 and R7 PRs. Pros (green) is expressed in two R7s. The dashed circle highlights an ommatidium (also see inset). Scale bar, 10µm. **c**, Ro (red) is transiently expressed near the morphogenetic furrow. Scale bar, 10µm. **d**, An ommatidium near morphogenetic furrow showing Ro expression in R2/5, and Bar expression in R3/4 and R1/6. Scale bar, 5µm. All nuclei are labeled with DAPI (gray).

**Extended Data Fig. 5.**
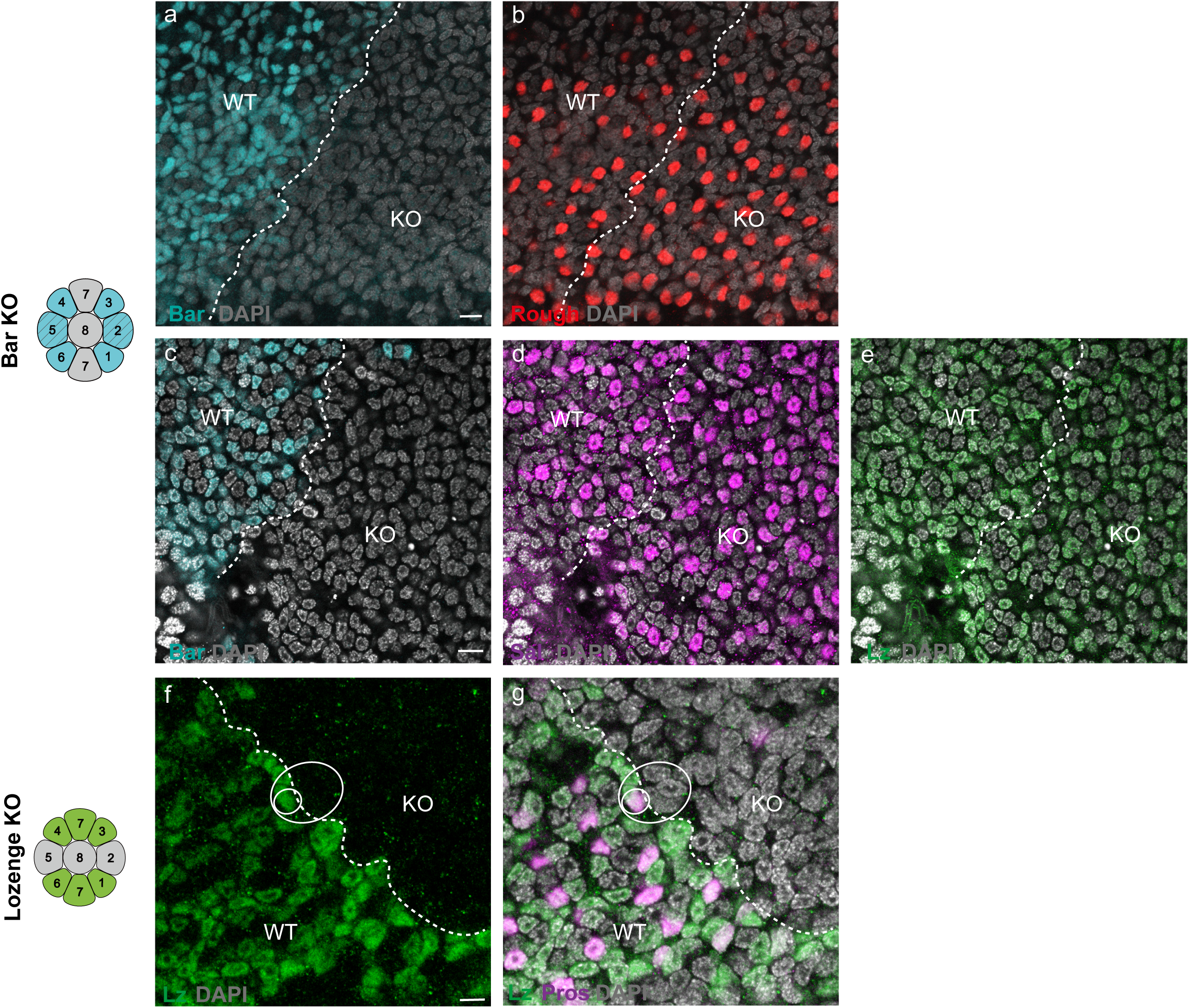
The effects of Bar and Lozenge loss-of-function in the butterfly *Vanessa* retina. **a-e**, CRISPR-induced mosaic knockout of *Bar* does not affect the expression of Ro (red, **b**), Sal (magenta, **d**) or Lz (green, **e**) in the pupal retina of *Vanessa*. Bar (cyan, **a**,**c**) is lost in KO regions compared to the WT region. The boundary between WT and Bar KO regions is indicated by a white dashed line. Scale bar, 5µm. **f-g**, CRISPR-induced mosaic knockout of *lz* results in the loss of R7s in the pupal retina of *Vanessa*. The solid white circle highlights an ommatidium near the clone boundary that has lost one R7 (the remaining R7 is stained with Lz (green, **f**) and Pros (magenta, **g**) on WT side) indicating that Lz is directly required in R7 recruitment. All nuclei are labeled with DAPI (gray). Scale bar, 10µm.

**Extended Data Fig. 6.**
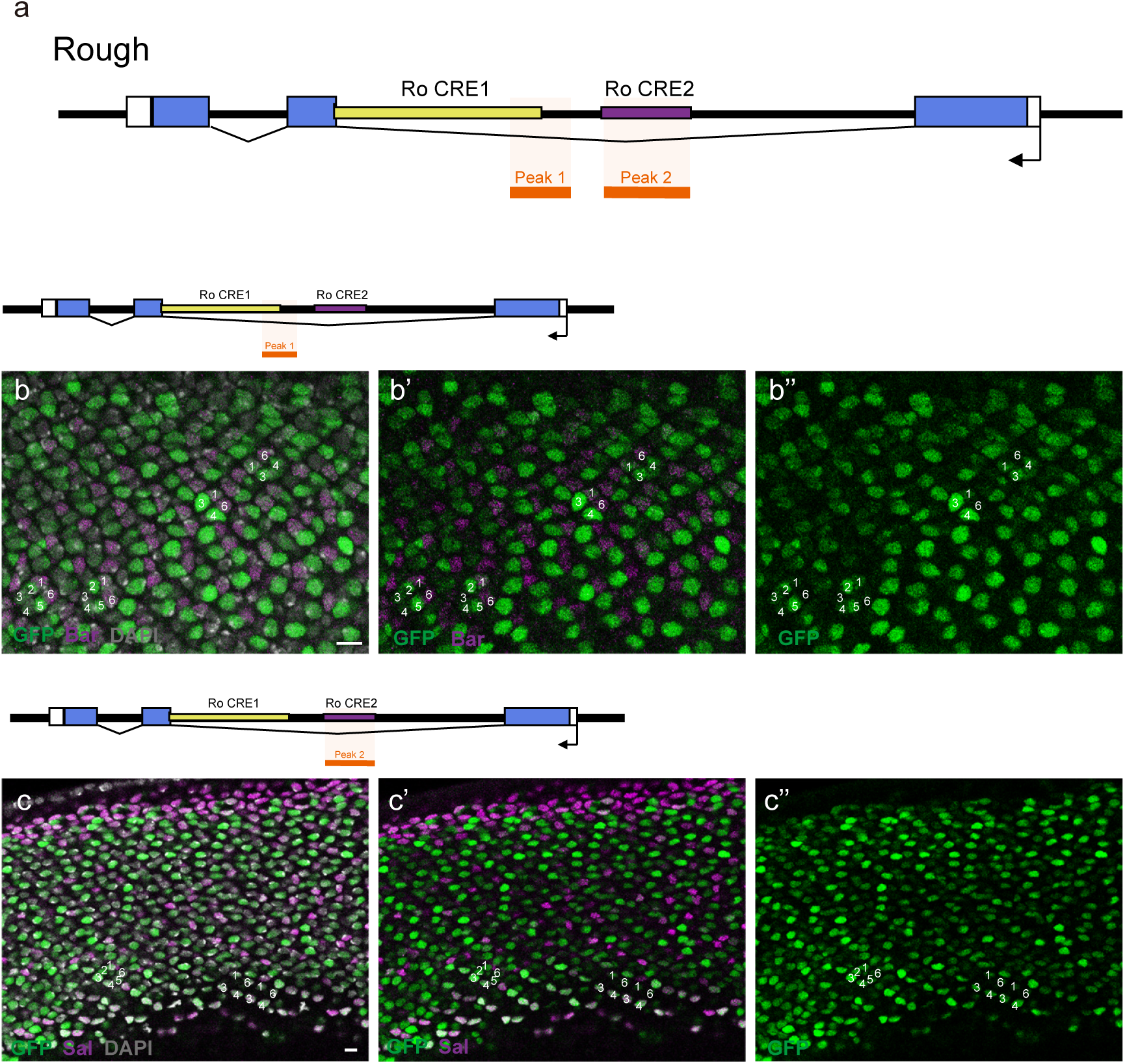
Rough CRE testing in *Drosophila*. **a**, Overview of the *Rough* gene in the *Drosophila* genome. Two Rough CREs were tested in intron 2, chosen partly based on single cell ATAC Seq data from^48^, with peaks shown in light orange. **b,** Ro CRE 1 driving expression of GFP (green) in the *Drosophila* 3^rd^ instar larval eye disc. Expression was variable in PRs 1-6, with Bar (magenta) expression shown in **b’** and GFP expression alone in **b’’**. **c,** Ro CRE 2 driving expression of GFP (green) in outer PRs 1-6, with Sal (magenta) labeling R3/4 in **c’.** GFP expression alone is shown in **c’’.** Scale bar, 5µm.

**Extended Data Fig. 7.**
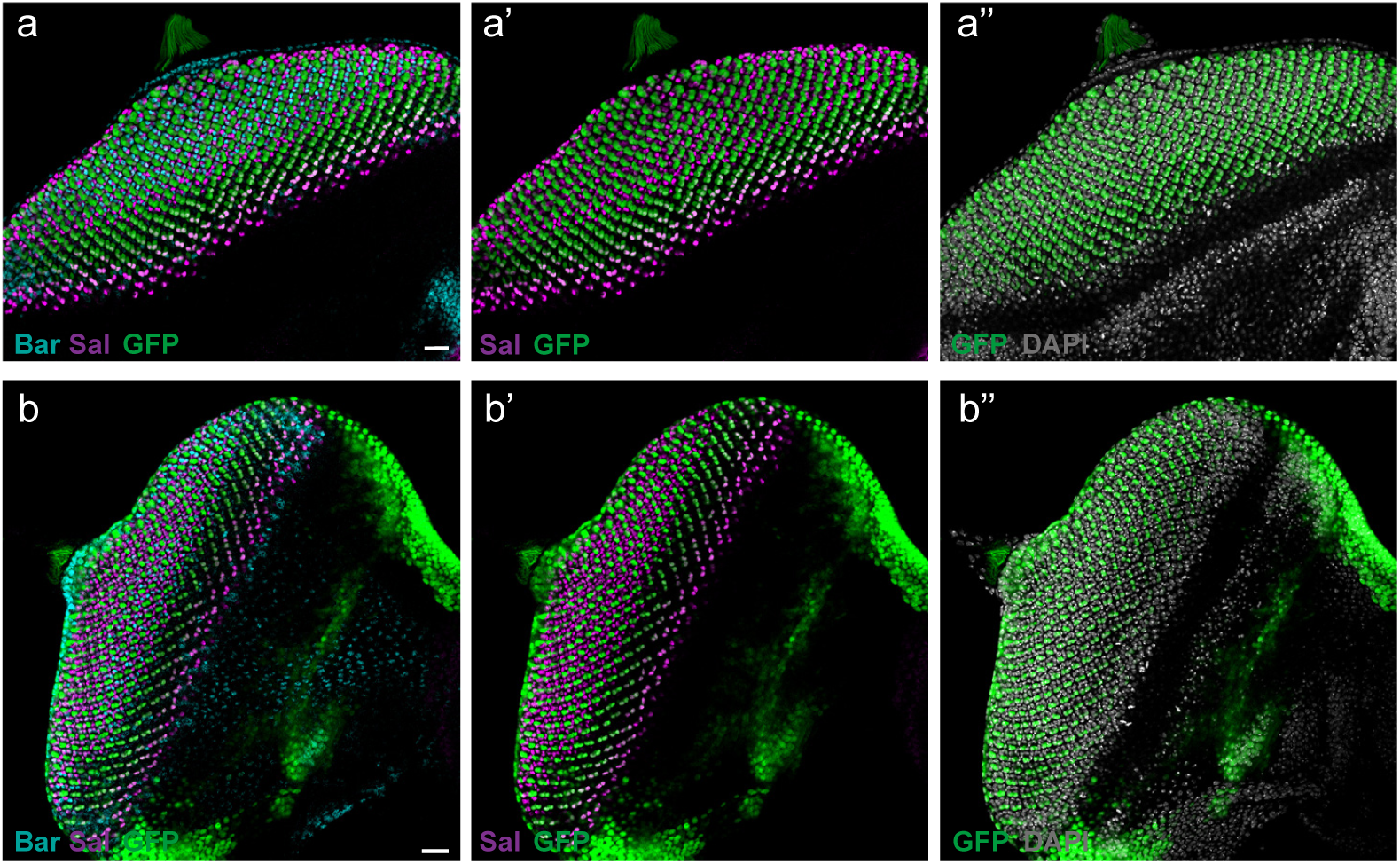
Two R3/4 CRE candidates in *Drosophila*. **a-a’’**, R55F01 from the CG14509 locus drive reporter expression in R2/5 and R3/4. GFP (green) labels R2/5 and R3/4. Sal (magenta) labels R3/4. Bar (cyan) labels R1/6. **b-b’’**, R86F02 from the Klumpfuss (Klu) locus drive reporter expression in R3/4. GFP (green) labels R3/4. Sal (magenta) labels R3/4. Bar (cyan) labels R1/6. All nuclei are labeled with DAPI (gray). Scale bar, 20µm.

**Extended Data Fig. 8.**
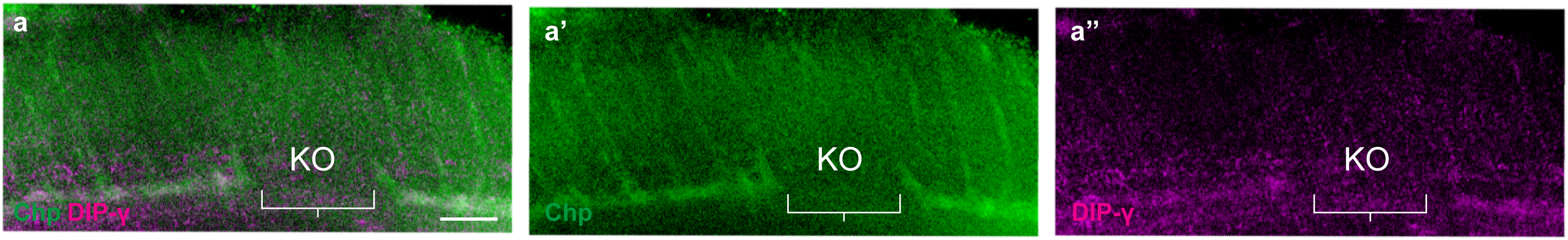
The effect of R7 loss on Dm8 retention in butterfly brains. **a**, CRISPR-induced mosaic knockout of *Lz* results in the loss of R7 axons and DIPγ expression in the *Vanessa* medulla. The *Lz* KO region (indicated by bracket) lacks Chp (green, **a’**) and DIPγ (magenta, **a’’**), which are visible in adjacent WT regions. Scale bar, 10µm.

**Extended Data Fig. 9.**
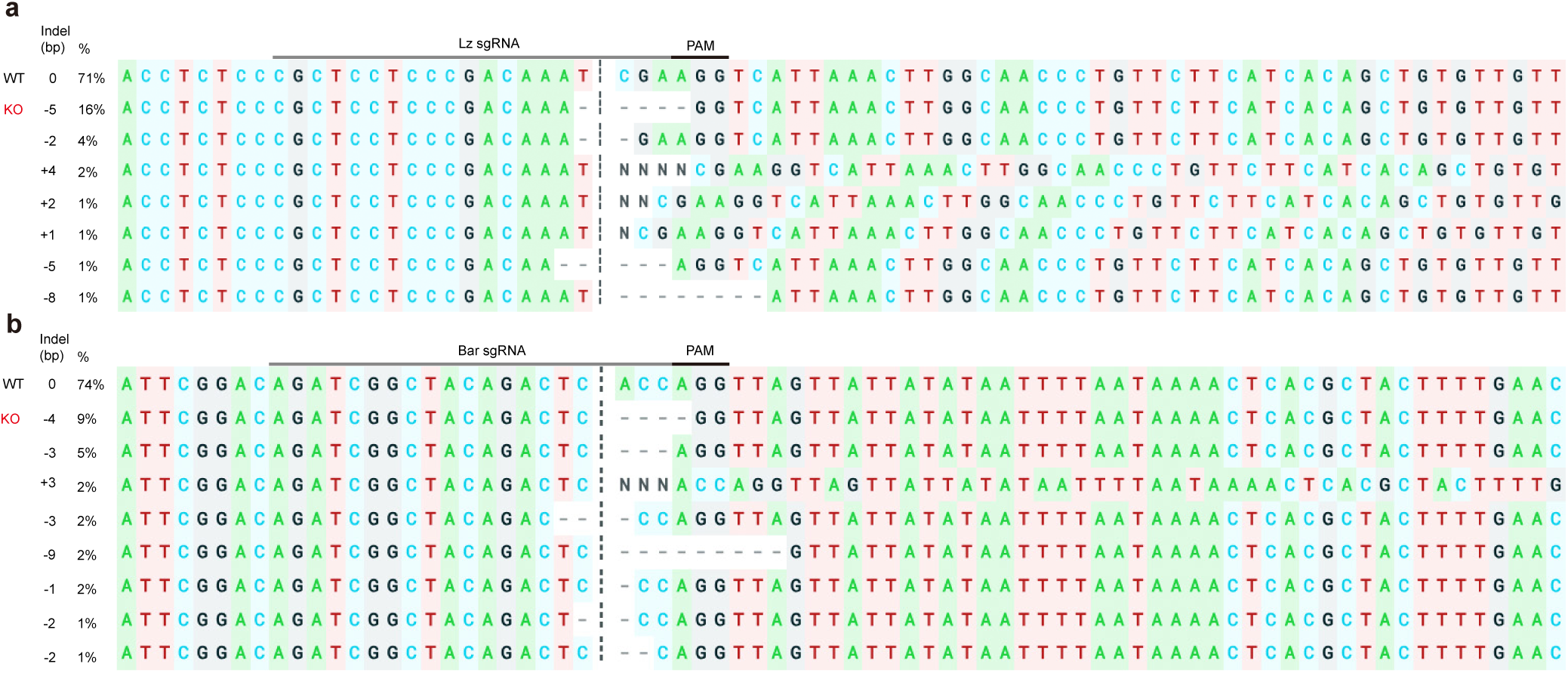
Genotyping validation of CRISPR-induced mosaic mutations. Sanger sequencing showing indels at the *Bar* (**a**) and *Lz* (**b**) target sites in *Vanessa* using Synthego ICE analysis. Predicted cut sites are indicated by dotted lines.

**Extended Data Fig. 10.**
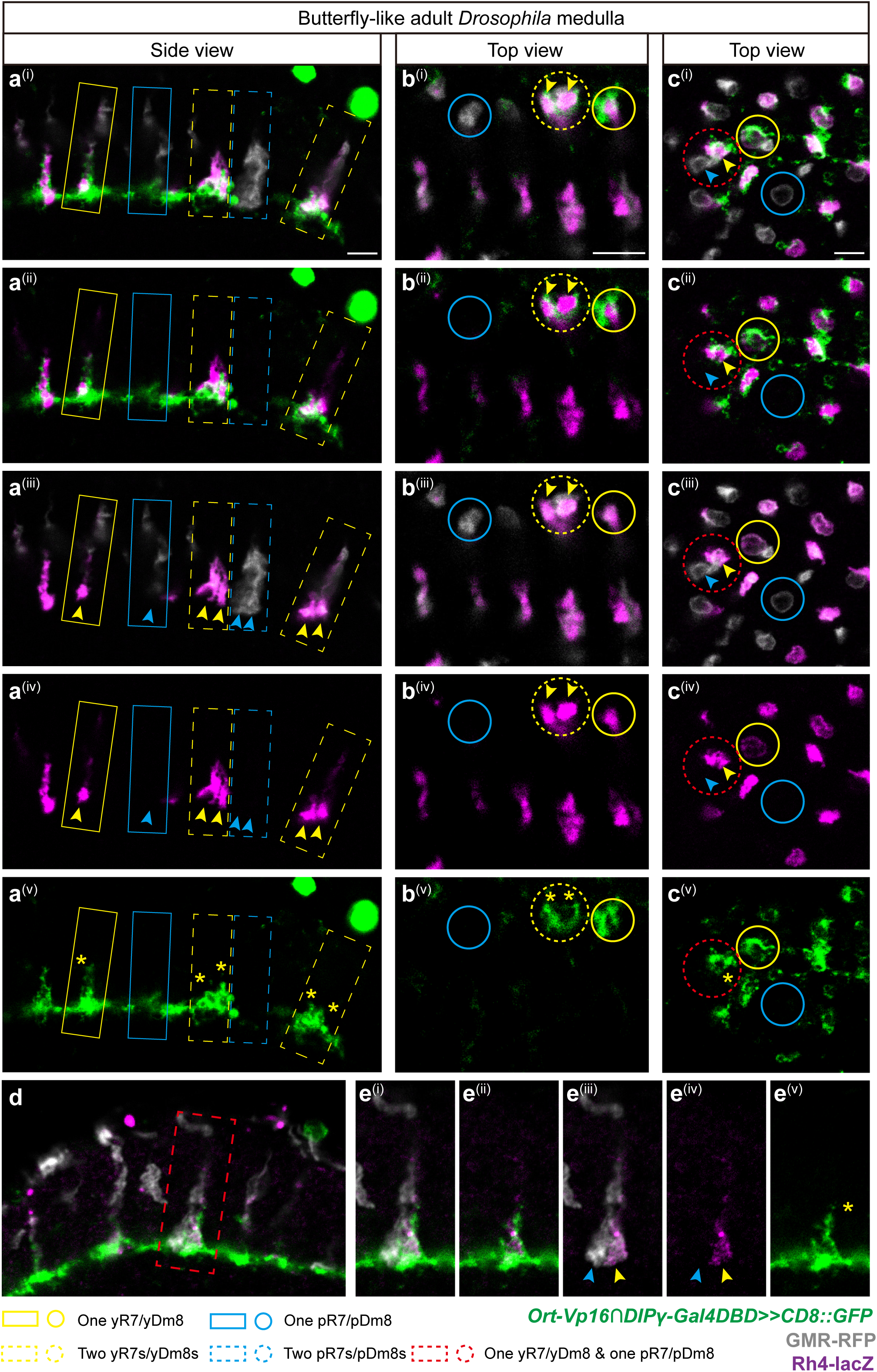
Specific connectivity between subtype R7 and Dm8 in butterfly-like adult *Drosophila* brain. **a^(i)^-a^(v)^**, side view of the adult medulla in sev.lz, sev.Su(H)enR flies. Comapred with WT columns containing one yR7 (yellow solid rectangle) and one pR7 (blue solid rectangle), yellow dashed rectangles highlight two yR7 axons (labelled by βGalactosidase, magenta) contacting two DIPγ-positive yDm8 neurons (green) within the same home column. Two yellow arrowheads indicate the two yR7 axons and two yellow asterisks mark the two DIPγ-positive yDm8 neurons in the same home column. Blue dashed rectangle highlights two pR7 axons (labelled by GMR-RFP, gray). Two blue arrowheads indicate the two pR7 axons and no DIPγ-positive yDm8 neurons are observed in the same home column. **b(i)-c(v)**, top-down view of the adult medulla in sev.lz, sev.Su(H)enR flies. Compared with WT columns containing one yR7 (yellow solid circle) and one pR7 (blue solid circle), yellow dashed circles in (**b^(i)^-b^(v)^**) highlight two yR7 terminals labelled by βGalactosidase (magenta, yellow arrowheads) that connect to two DIPγ-positive yDm8 neurons (green, yellow asterisks) within a single column. Red dashed circles in (**c^(i)^-c^(v)^**) highlight two R7 terminals, one yR7 labelled by βGalactosidase (magenta, yellow arrowhead) and one pR7 labelled by GMR-RFP (gray, blue arrowhead) within a single home column, in which one DIPγ-positive yDm8 neuron (green) connects to the yR7 terminal and no DIPγ-positive yDm8 neuron contacts the pR7 terminal. **d-e^(v)^**, side view of the adult medulla in sev.lz, sev.Su(H)enR flies. The red dashed rectangle in (**d**) highlights two R7 axons, one yR7 axon labelled by βGalactosidase (magenta, yellow arrowhead) and one pR7 axon labelled by GMR-RFP (gray, blue arrowhead) within a single home column, in which one DIPγ-positive yDm8 neuron (green, yellow asterisk) connects to the yR7 axon and no DIPγ-positive yDm8 neuron contacts the pR7 axon. Scale bar, 5µm.

**Supplementary Table 1:**
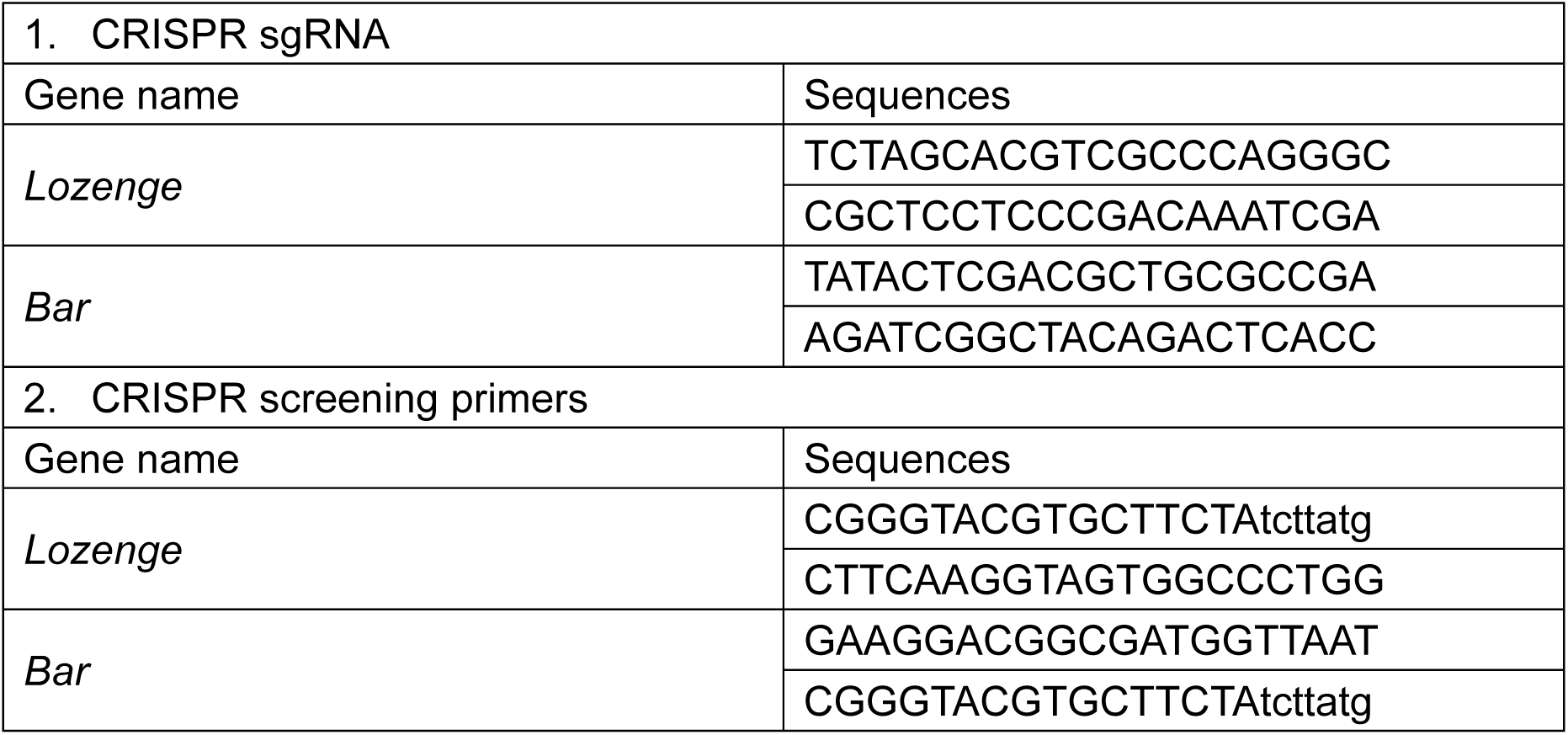
Primers and sgRNA guides used in this study.

**Supplementary Table 2:**
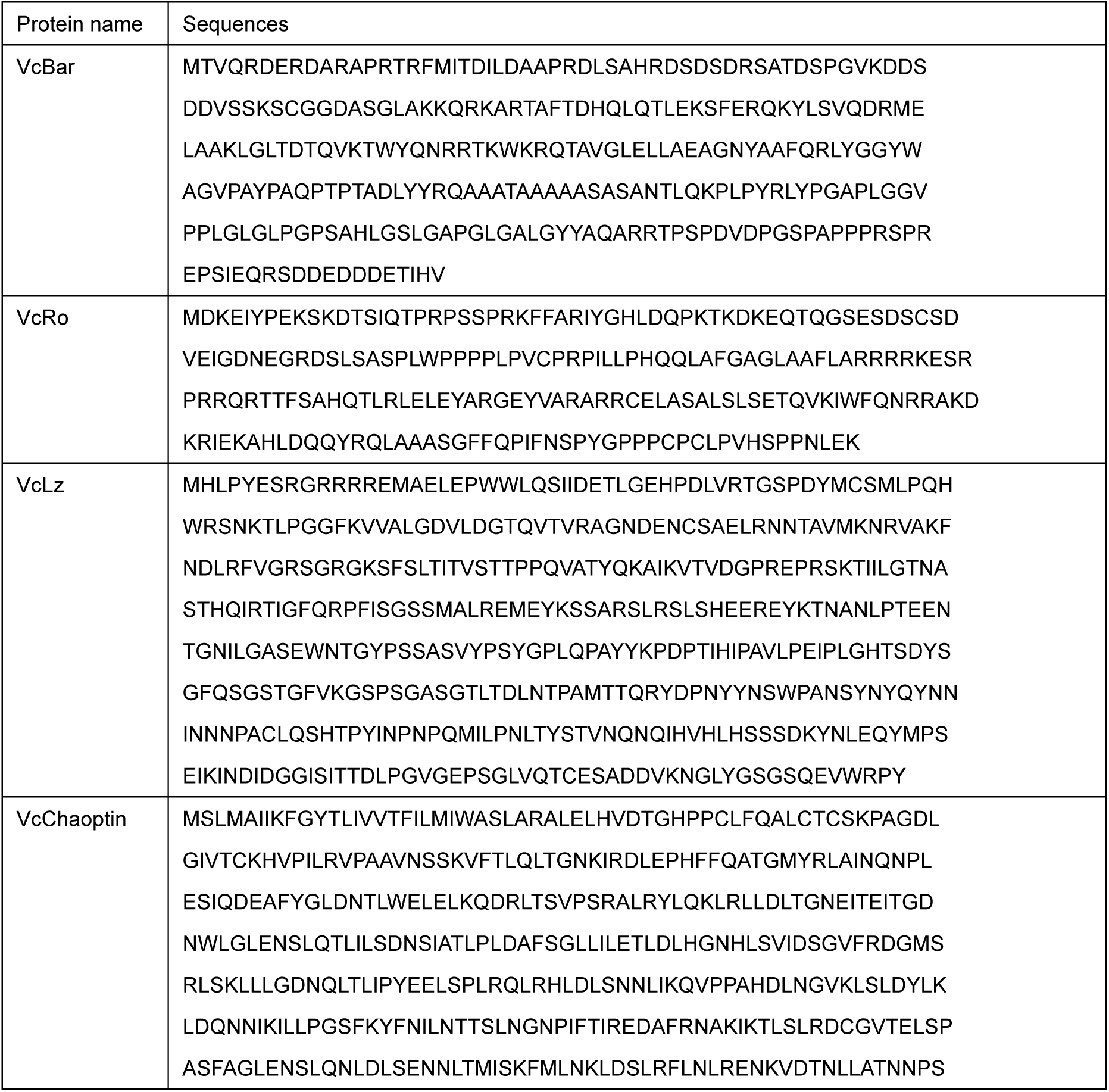
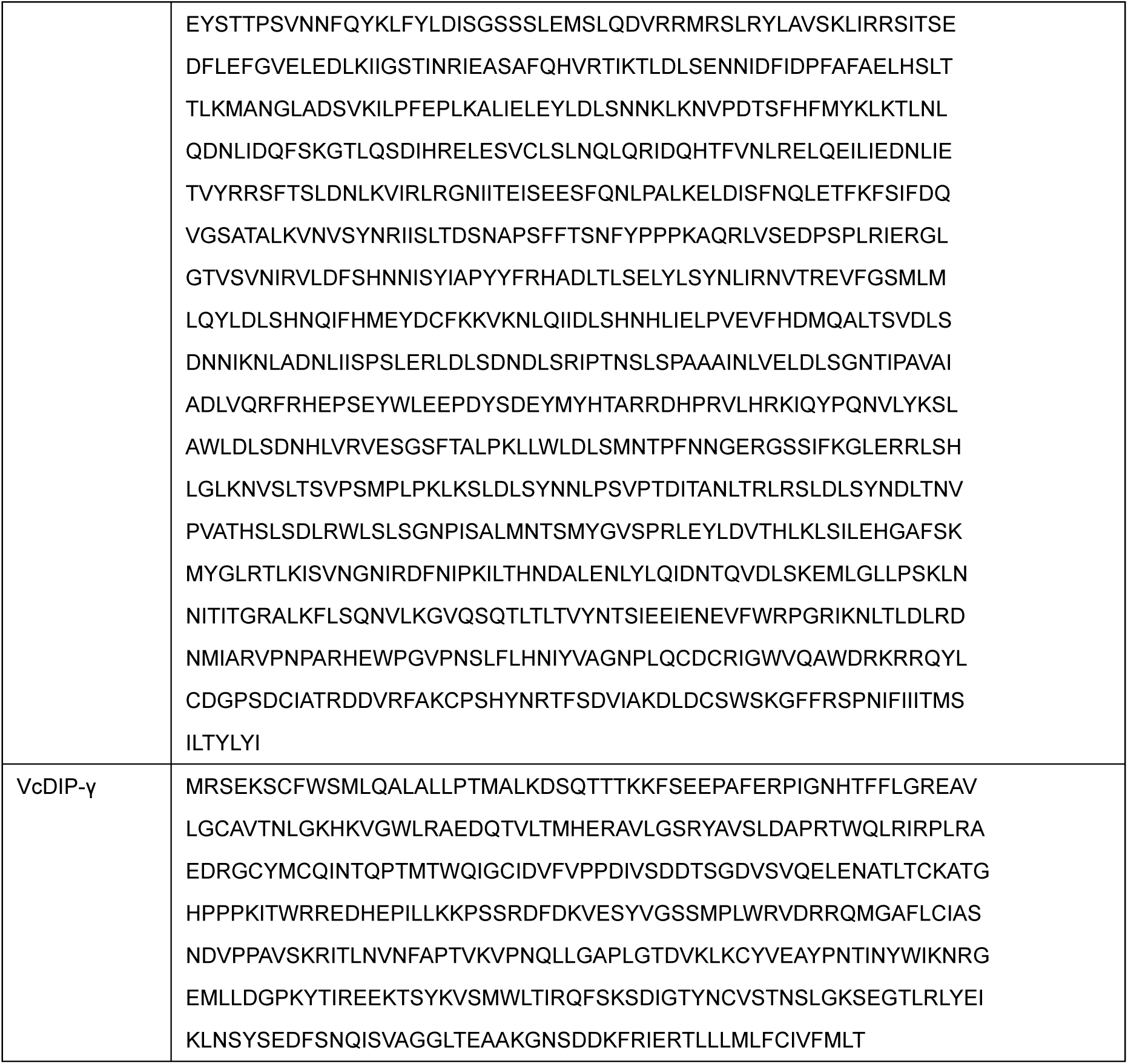
Sequences used to produce antibody.

**Supplementary Table 3:**
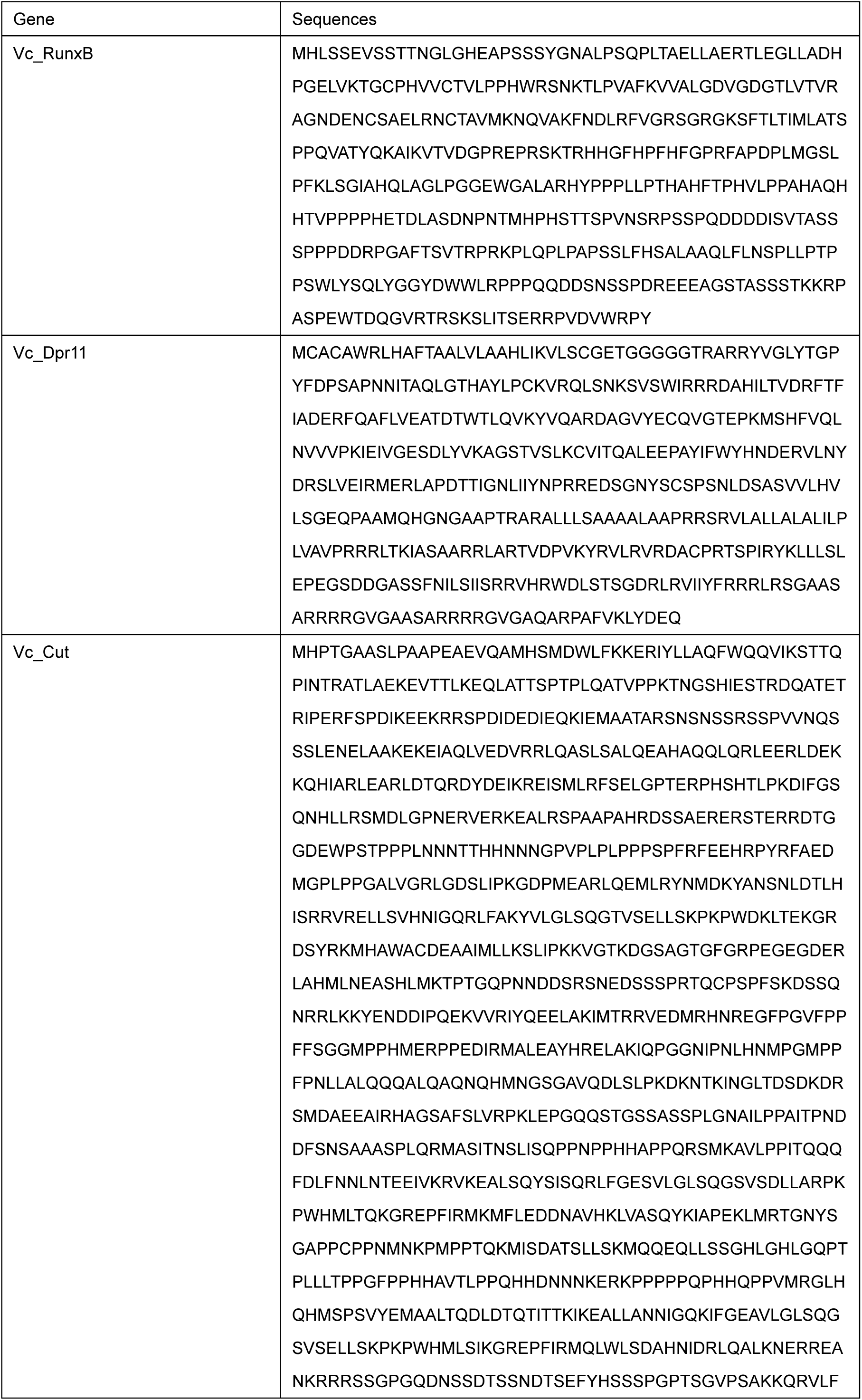
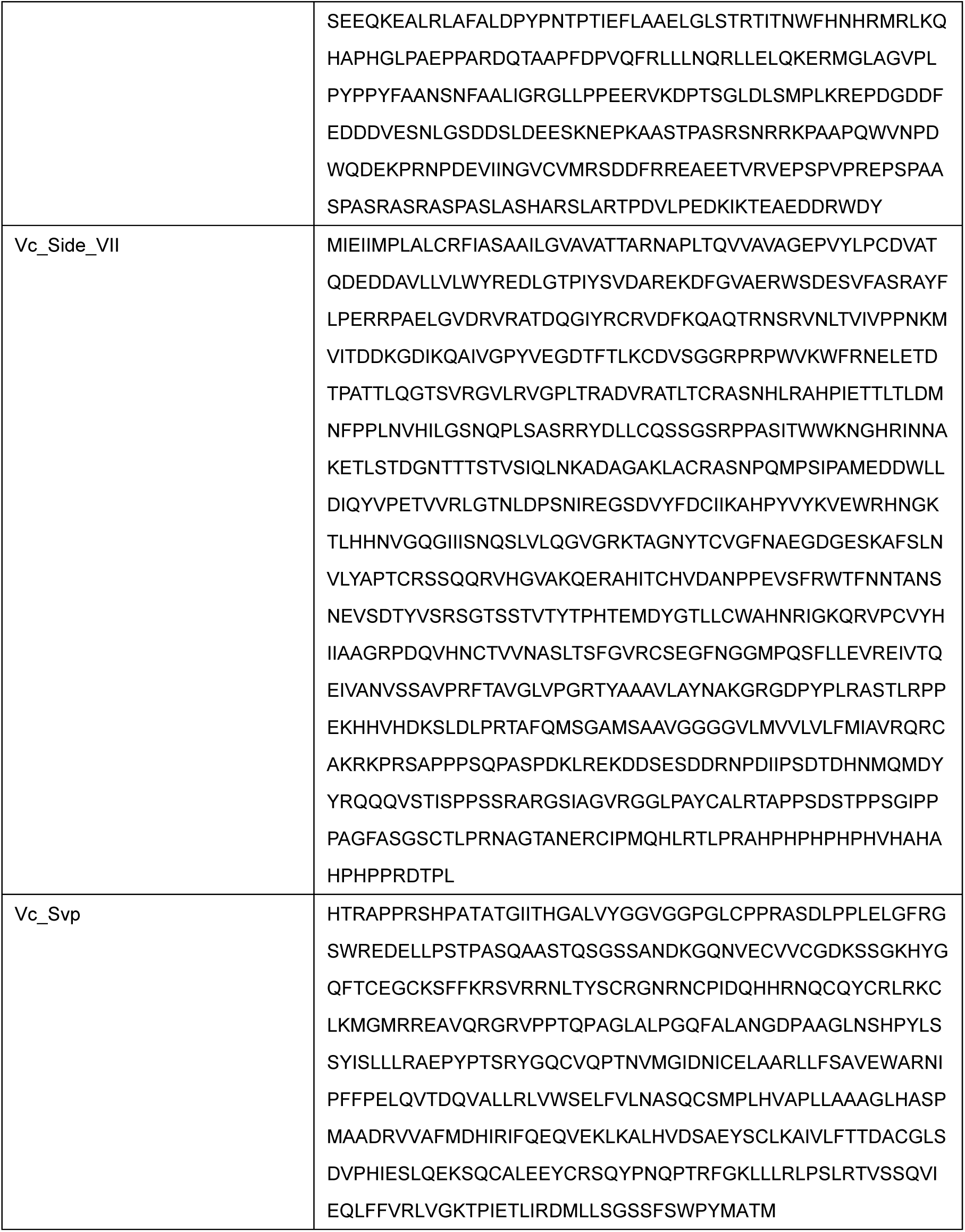
Sequences used to design HCR probes.

**Supplementary Table 4:**
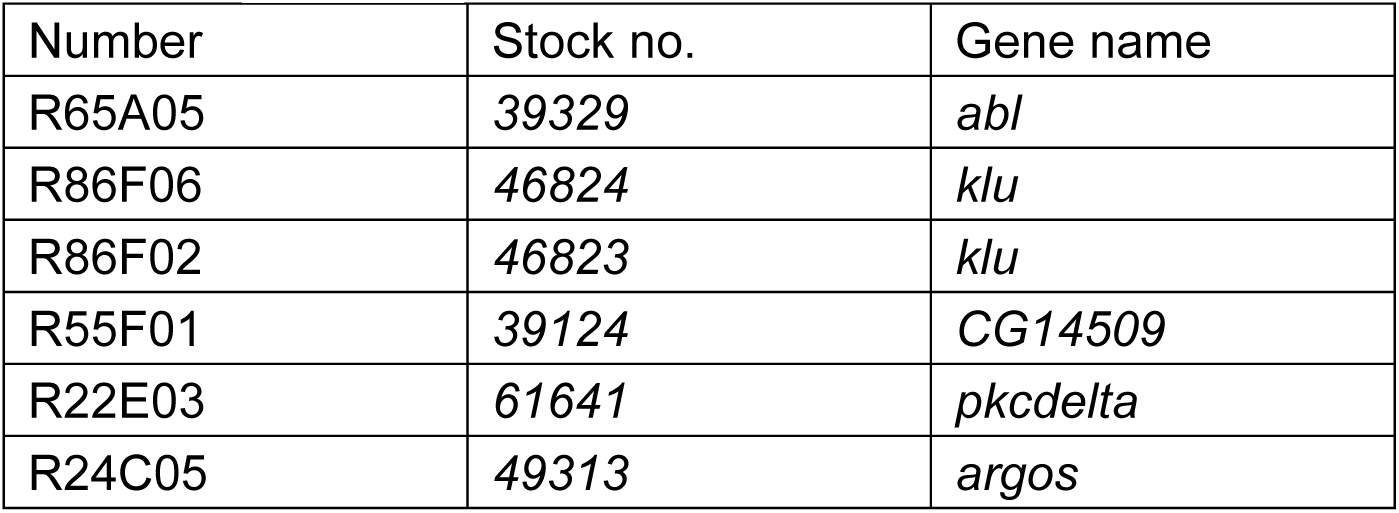
Six of the 61 lines from the Janelia FlyLight collection that were evaluated as potential R3/4 enhancers.

## Notes

### Competing Interest Statement

The authors have declared no competing interest.

### Summary of Updates

This version of the manuscript has been revised to add new data. We added a new Figure 6 where we examined subtype specificity of connections of R7 photoreceptors in flies that produce butterfly-like ommatidia.

